# A high-resolution protein-RNA binding atlas for structure-aware interaction prediction and post-transcriptional regulatory modeling

**DOI:** 10.64898/2026.10.12.758580

**Authors:** Mehran Karimzadeh, Lisa Fish, Heather Karner, Johannes Linder, Ali Emadi, Aldo Hernandez-Corchado, Fatima Zohra Smaili, Khashayar Ghavami, Edna S. Stewart, David Kelley, Bo Wang, Hamed Shateri Najafabadi, Manuel D. Leonetti, Hani Goodarzi

**Author notes:** Corresponding authors: David Kelley, Bo Wang, Hamed Shateri Najafabadi, Manuel Leonetti, Hani Goodarzi. _These authors contributed equally to this work._ (Mehran Karimzadeh, Lisa Fish, Heather Karner, Johannes Linder, Ali Emadi).

## Abstract

RNA-binding proteins (RBPs) control post-transcriptional gene expression by regulating RNA processing, localization, stability, and translation. However, our understanding of RNA regulation remains limited by the resolution, scale, and uniformity of available RBP binding maps. Here we introduce eclair CLIP (ecCLIP), a high-throughput CLIP-seq platform that combines endogenous protein tagging for uniform affinity capture with ligation-free library preparation to generate high-resolution RNA-protein binding profiles. We applied ecCLIP to 67 endogenously tagged RBPs in HEK293T cells, generating a high-quality transcriptome-scale atlas that recovered known binding preferences and captured post-transcriptional regulons linked to RNA half-life. To test the utility of the atlas, we incorporated ecCLIP into Borzuki, a Borzoi-derived sequence-to-function model trained on complementary RNA degradation, expression, CLIP, and Perturb-seq data. Inclusion of ecCLIP data improved RNA half-life prediction and assessment of 3^′^ UTR variant effects on RNA stability, demonstrating that our atlas helps capture RNA dynamics. To directly model the cis-regulatory grammar underlying these RBP-RNA interactions, we developed Pythia, a structure-aware neural network architecture that incorporates context-free grammar-inspired fixed convolutions to explicitly encode RNA base-pairing features. Pythia predicted RBP binding from sequence and structural context and decomposed predictions into sequence- and structure-dependent components. We tested this decomposition for five RBPs using massively parallel reporter CLIP (MPRC) binding assays in cells, using synthetic RNA libraries containing mutations designed to preserve or disrupt predicted RNA structures across nominated binding sites. We used the same MPRC strategy to confirm allele-specific binding effects for variants prioritized by Pythia. Together, ecCLIP and Pythia provide a resource for structure-aware decoding of RNA-protein interaction grammar and interpretation of post-transcriptional regulatory variation.

## 1. Introduction

RNA-binding proteins (RBPs) regulate gene expression after transcription by controlling RNA processing, localization, stability, and translation. These interactions define post-transcriptional regulatory programs that determine when, where, and how transcripts are expressed. Disruption of RBP abundance, binding specificity, or cis-regulatory RNA elements that they recognize contributes to diverse human diseases (Gerstberger et al., 2014; Hentze et al., 2018; Gebauer et al., 2021). Understanding and predicting these mechanisms requires transcriptome-scale maps of RBP binding and models that connect binding events to RNA fate.

Crosslinking immunoprecipitation followed by sequencing (CLIP-seq) is a central approach for mapping RBP-RNA interactions in cells (Ule et al., 2003; Licatalosi et al., 2008; König et al., 2010; Hafner et al., 2010; Van Nostrand et al., 2016, 2020). Existing CLIP resources have provided valuable maps of RBP occupancy, but high-quality profiling across many RBPs remains technically demanding. Antibody-specific immunoprecipitation conditions, radioactive visualization, ligation-dependent library preparation, and protocol heterogeneity limit throughput, consistency, and recovery. These limitations are especially consequential for sequence-to-function modeling, where each experimental track must be sufficiently resolved, uniform, and biologically informative to improve generalization.

To address this need, we developed eclair CLIP (ecCLIP), a high-throughput CLIP-seq platform for systematic RBP profiling that combines endogenous split-fluorescent-protein tagging for uniform affinity capture with infrared visualization and ligation-free library preparation. We applied ecCLIP to 67 endogenously tagged RBPs in HEK293T cells, generating a high-resolution transcriptome-scale atlas that captures RBP binding specificity and regulatory signal linked to RNA half-life. We reasoned that a high quality RBP binding atlas should measurably improve our ability to decode post-transcriptional regulatory grammars beyond what existing public binding and RNA processing datasets allow. We built Borzuki to test this directly. Borzuki is a Borzoi-derived (Linder et al., 2025) sequence-to-function model for RNA expression and dynamics that extends the sequence-to-stability framing introduced by Saluki (Agarwal and Kelley, 2022). It takes ecCLIP profiles for stability-modulating RBPs as its central new input, alongside complementary and publicly available ENCODE RBP eCLIP (Van Nostrand et al., 2020), RBP Perturb-seq (Kowalski et al., 2024), and RNA decay data. In matched ablations, adding ecCLIP data improved prediction of RNA half-life and of MPRA-measured 3^′^UTR variant effects on RNA stability, showing that the ecCLIP atlas carries regulatory information about RNA fate that existing data does not.

Beyond their utility for predicting RNA regulation, nucleotide-resolution RNA interaction profiles also provide an opportunity to learn the underlying *cis*-regulatory grammars. RBP-RNA binding is shaped not only by primary sequence but also by RNA base-pairing, structural accessibility, and higher-order structural context (Eddy and Durbin, 1994; Dowell and Eddy, 2004; Lange et al., 2012; Goodarzi et al., 2014; Kazan et al., 2010). Earlier work has shown that structural RNA regulatory elements can encode post-transcriptional programs (Goodarzi et al., 2014, 2012). However, most deep learning models of RBP binding represent RNA sequence explicitly while treating structure implicitly, if at all.

To address this we developed Pythia, a structure-aware neural network architecture that explicitly models the cis-regulatory grammar of RBP-RNA recognition. Pythia encodes RNA base-pairing features through context-free grammar-inspired fixed convolutions while preserving a dedicated sequence arm for primary sequence recognition, enabling both prediction of RBP binding and decomposition of individual binding sites into sequence- and structure-dependent components. To experimentally validate these predictions, we developed massively parallel reporter CLIP (MPRC), the first approach enabling pooled, quantitative measurement of sequence- and structure-dependent binding effects alongside allele-specific effects for prioritized regulatory variants.

This study is organized in three parts. First, we describe the generation of the ecCLIP atlas and characterize its quality and resolution. Second, we evaluate its functional utility through Borzuki, demonstrating that ecCLIP profiles improve prediction of RNA half-life and 3^′^ UTR variant effects beyond what existing data afford. Third, we deploy Pythia to dissect the sequence and structural determinants of RBP recognition and interpret the binding consequences of regulatory variants. Together, these results establish an integrated framework in which high-resolution RBP binding maps sharpen models of RNA fate and enable mechanistic interpretation of RNA-protein interaction grammar at nucleotide resolution.

## 2. Results

### 2.1. Generating a high-resolution RBP binding atlas with eclair CLIP

CLIP-seq remains a central technique for transcriptome-wide mapping of RBP-RNA interactions, but applying CLIP systematically across large RBP panels is limited by several practical and technical bottlenecks. Conventional workflows typically require protein-specific antibodies and immunoprecipitation conditions, and differences in antibody availability, specificity, affinity, and lot-to-lot quality make atlas-level comparisons across RBPs difficult. Other steps, especially ligation-dependent library preparation, can reduce recovery and introduce bias. CLIP signal is also influenced by crosslinking efficiency, which can complicate interpretation of broad peaks and lower-resolution binding profiles (Park et al., 2018). Subsequent methods have addressed subsets of these limitations, including size-matched input controls to account for background abundance (Van Nostrand et al., 2016) and photoactivatable ribonucleoside labeling to identify binding sites through diagnostic mutational signatures (Ascano et al., 2012). TAG-eCLIP further addressed dependence on RBP-specific antibodies by combining CRISPR-mediated endogenous epitope tagging with tag-specific immunoprecipitation (Van Nostrand et al., 2017). However, a uniform and efficient workflow is still needed to generate high-quality, high-resolution RBP binding maps at scale.

To enable systematic profiling of RBP binding, we developed eclair CLIP (ecCLIP), an efficient and streamlined CLIP-seq workflow optimized for high-throughput, high-resolution mapping of ribonucleoprotein complexes (**Fig. 1a**). ecCLIP combines endogenous split mNeonGreen (mNG) tagging for uniform affinity capture, infrared labeling in place of radioisotope-based visualization, and ligation-free, template-switch-based library preparation to improve recovery and reduce ligation-associated bias (Feng et al., 2017; Zarnegar et al., 2016). We evaluated recovery of mNG-tagged RBPs, optimized RNA labeling, and compared RNA isolation methods (**Supp. Fig. S1a–e**).

**Figure 1.**
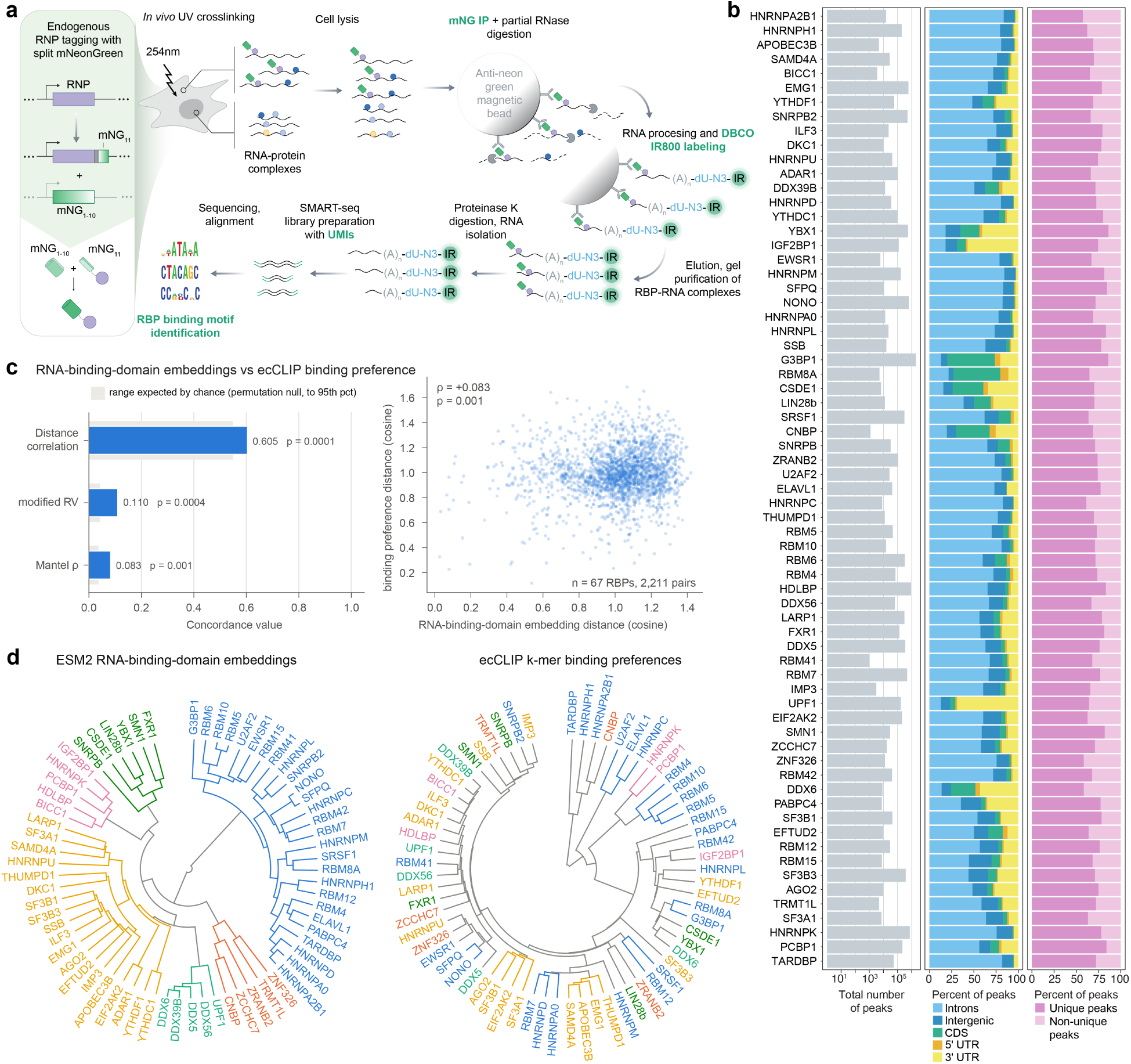
Efficient, high-resolution profiling of RBP-RNA interactions with eclair CLIP. **(a)** Schematic of the ecCLIP workflow. Endogenous split mNeonGreen tagging provides a uniform affinity-capture strategy across RBPs. After UV crosslinking, cell lysis, partial RNase digestion, anti-mNG immunoprecipitation, infrared RNP labeling, gel separation, proteinase K digestion, RNA isolation, and ligation-free template-switch library preparation with UMIs, sequencing reads are aligned to identify transcriptome-wide RBP binding profiles. **(b)** Barplots show the number of peaks detected per RBP (left), the fraction of peaks overlapping UTRs, coding sequence, intergenic regions, and introns (middle), and the fraction of peaks that are unique or shared across RBPs (right). **(c)** Correspondence between ESM2 RNA-binding-domain embeddings and ecCLIP-derived binding preferences for 67 RBPs. Left, concordance statistics (distance correlation, modified RV coefficient, and Mantel *ρ*); grey bands show the range expected by chance (95th percentile of 9,999 permutations of RBP labels). Right, pairwise cosine distances between RBP domain embeddings versus between binding-preference profiles (2,211 RBP pairs). **(d)** Circular dendrograms of the 67 RBPs clustered (Ward, k = 6) by ESM2 RNA-binding-domain embeddings (left) and by ecCLIP-derived 5-mer binding preferences (right). Both trees are colored by the same grouping, defined from the protein-embedding clusters, so scattered colors in the right tree show where the two representations disagree.

We selected RBPs primarily from the OpenCell endogenous tagging resource (Cho et al., 2022) and supplemented this set with additional proteins to represent major RBP families. The final panel included RBPs containing a wide variety of RNA-binding domains, including RNA recognition motifs, zinc finger motifs, KH-domains, DEAD-box helicases, YTH domains and double-stranded RNA-binding domains (**Supp. Fig. S1f**). We also designed the atlas to complement the ENCODE eCLIP resource in two ways: by profiling RBP binding in HEK293T cells, rather than the K562 and HepG2 cell contexts used for most ENCODE eCLIP datasets, and by including 40 RBPs not represented in ENCODE. In total, we profiled 67 endogenously tagged RBPs and defined their transcriptome-wide binding landscapes in HEK293T cells.

Across all 67 RBPs, we identified 9,097,672 ecCLIP peaks (**Supp. Fig. S1g**). Peak counts varied substantially across proteins, which can be attributed to a variety of factors, including RBP-specific differences in expression, crosslinking efficiency, RNA occupancy, and biochemical recovery (**Fig. 1b**, left). The resulting binding profiles also showed distinct regional preferences, with examples of RBPs enriched in UTRs (ELAVL1), coding sequences (YBX1), or intronic regions (HNRNPC) (**Fig. 1b**, middle). Most peaks were unique to individual RBPs, whereas a relative minority was shared between two or more RBPs (**Fig. 1b**, right), consistent with the atlas capturing largely protein-specific binding programs.

We compared pairwise distances between ESM2 protein embeddings and ecCLIP-derived binding preferences to assess correspondence between these representations (Ray et al., 2013). Here, binding preferences were represented as dense 5-mer enrichment profiles of ecCLIP peaks relative to flanking background, a compact description of each RBP’s RNA sequence specificity, and proteins were represented by ESM2 embeddings of their annotated RNA-binding domains. Distances in the two representations showed limited but non-random correspondence (Mantel *ρ* = 0.083, modified RV coefficient = 0.110, distance correlation = 0.605; each exceeding its permutation null, P ≤ 0.002; **Fig. 1c**), and clustering the 67 proteins by domain embeddings and by ecCLIP-derived sequence preferences produced largely different groupings (**Fig. 1d**). Whole-protein embeddings gave a similar, slightly stronger correspondence (Mantel *ρ* = 0.117, modified RV = 0.121, distance correlation = 0.641; P = 0.0001; **Supp. Fig. S1i,j**). This comparison does not establish the limits of motif prediction from protein sequence: RNA-binding-domain similarity and models trained specifically to predict RNA preferences address that question more directly. Rather, it shows that distances between general-purpose ESM2 embeddings, which are not trained on RNA-binding data, only weakly reflect the in vivo sequence preferences measured by ecCLIP, and therefore should not be used on their own as a proxy for shared binding specificity (Ray et al., 2013; Sasse et al., 2026). These results establish ecCLIP as a high-resolution atlas for empirically measuring RBP binding programs across a broad panel of proteins.

### 2.2. Establishing ecCLIP data quality and utility for modeling RNA dynamics

We next evaluated the resolution and signal quality of the ecCLIP atlas using two complementary metrics. First, we quantified the fraction of reads in peaks (FRiP) across window sizes centered on peak summits, an analysis that measures how tightly the sequencing signal is concentrated around inferred binding sites. For nine RBPs (TARDBP, HNRNPK, HNRNPC, PCBP1, EWSR1, HNRNPL, IGF2BP1, LIN28B, and ZRANB2), peak-centered FRiP curves and aggregate read profiles showed concentrated ecCLIP signal around inferred binding sites (**Fig. 2a**; **Supp. Fig. S2a**). Published eCLIP profiles are included as a reference rather than as a controlled comparison of assay performance, because they differ in cellular context, library preparation, and peak definition (Van Nostrand et al., 2020). Second, to test whether ecCLIP peaks capture biologically meaningful RBP binding preferences, we compared sequence preferences inferred from cell-derived ecCLIP peaks to orthogonal in vitro measurements from RNA Bind-n-Seq (RBNS; Lambert et al., 2014). For TARDBP (TDP-43), ecCLIP recovered GAAUG-containing 5-mers consistent with its RNAcompete consensus motif, providing an example of recognizable sequence specificity (**Fig. 2b**; Ray et al., 2013). We then assessed concordance with RBNS across up to nine RBPs with available eCLIP, ecCLIP, and RBNS measurements. For each RBP, we computed the absolute Pearson correlation between CLIP-derived k-mer enrichment and RBNS affinity at the highest reported RBNS protein concentration (1300 nM), using k = 4, 5, and 6 and peak-centered windows of 10 or 20 nucleotides. ecCLIP-derived k-mer enrichments agreed with RBNS preferences across these parameter settings; the corresponding eCLIP measurements provide a contextual reference (**Fig. 2c**). This result was robust when RBNS affinity was evaluated at the optimal protein concentration for each RBP rather than at a fixed concentration (**Supp. Fig. S2b**). Together, these results show that ecCLIP produces binding maps that are both high resolution and informative of RBP sequence specificity.

**Figure 2.**
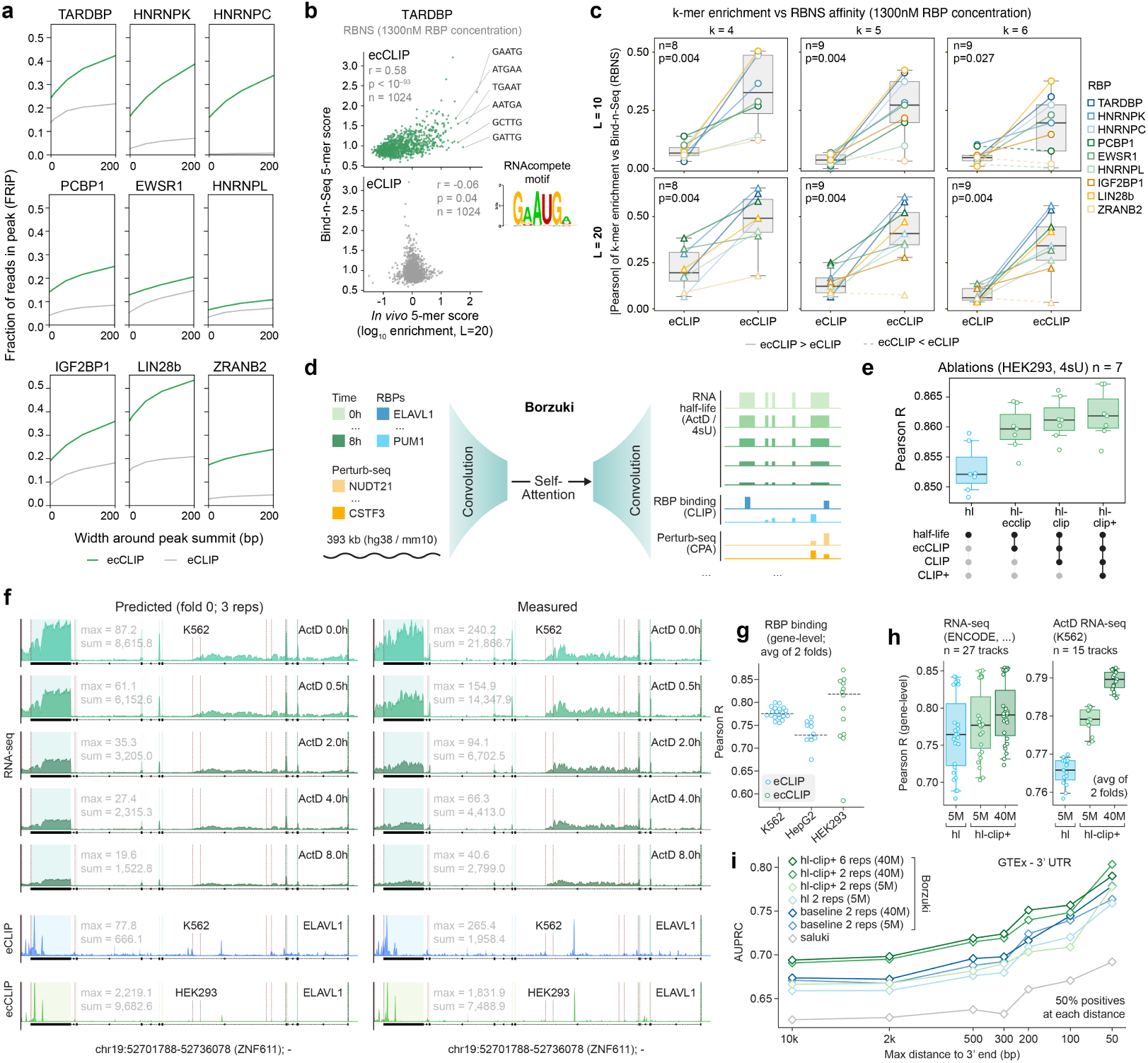
ecCLIP quality metrics and genome-scale modeling of RNA dynamics. **(a)** Comparison of peak resolution in ecCLIP and eCLIP. Fraction of reads in peaks (FRiP) is shown as a function of window width around each peak summit for nine RBPs with comparable datasets (TARDBP, HNRNPK, HNRNPC, PCBP1, EWSR1, HNRNPL, IGF2BP1, LIN28B, and ZRANB2). **(b)** Example comparison between CLIP-derived and RBNS-derived sequence preferences for TARDBP. Scatterplots compare 5-mer enrichment scores (log ratio of peak vs flanking background, 20-nt peak-centered windows) from eCLIP or ecCLIP peaks to RBNS 5-mer affinity scores at 1300 nM. The top enriched 5-mers are labeled, and the TDP-43 RNAcompete motif is shown for reference. **(c)** Systematic comparison of k-mer concordance between eCLIP/ecCLIP and RBNS across up to nine RBPs. The absolute Pearson correlation between CLIP-derived k-mer enrichment and RBNS affinity at 1300 nM is shown for k = 4, 5, and 6, using peak-centered windows of 10 or 20 nucleotides. Lines connect matched RBPs between eCLIP and ecCLIP; solid lines indicate stronger concordance for ecCLIP and dashed lines indicate stronger concordance for eCLIP. P-values correspond to paired comparisons between eCLIP and ecCLIP. **(d)** Borzuki training data and architecture overview. Borzuki is trained to jointly predict sequencing coverage from RNA-seq degradation time courses, RBP CLIP-seq, and 3^′^-processing regulator Perturb-seq, together with ENCODE RNA-seq/DNase-seq and FANTOM5 CAGE data. The 40-million-parameter transformer U-net predicts coverage profiles for 452 human and 160 mouse tracks from 393-kb input sequences and is trained as an ensemble of six replicate models. **(e)** Borzoi-style model ablation for HEK293T RNA half-life prediction. Boxplots summarize Pearson correlations (n = 7 RNA-seq tracks) for models trained with half-life tracks (‘hl’), half-life tracks plus ecCLIP (‘hl_ecclip’), half-life tracks plus ecCLIP and eCLIP (‘hl_clip’), or half-life tracks plus both CLIP resources and 3^′^-end processing Perturb-seq (‘hl_clip+’). Reported metrics were averaged across 2 model replicates. **(f)** Predicted and measured coverage for the held-out gene ZNF611 using a three-model ensemble. Top: RNA-seq coverage across K562 actinomycin D time points (0-8 h). Bottom: ELAVL1 binding from two independent experiments, ENCODE eCLIP in K562 and ecCLIP in HEK293T, shown as representative CLIP-seq tracks predicted by the same model. **(g)** Test-set Pearson correlation for CLIP-seq gene-level prediction across held-out tracks. **(h)** Model size and training-data composition comparison. Pearson correlation metrics are shown for gene-level log-coverage prediction across two model sizes (5 million and 40 million parameters) and two training-data compositions, with or without CLIP-seq and Perturb-seq modalities. **(i)** Comparison between Saluki and Borzuki at distinguishing fine-mapped 3^′^ UTR eQTLs (PIP > 0.7) from matched negative SNPs (Kerimov et al., 2021; Lonsdale et al., 2013). Borzuki model size, ensemble size, and training-data composition are varied.

For this evaluation of data utility, we developed Borzuki as an expansion of Borzoi focused on post-transcriptional regulation (“Borzuki” being a combination of Borzoi and Saluki; Agarwal and Kelley, 2022). Borzoi established that long-range RNA-seq coverage can be predicted directly from genomic sequence, providing a general framework for learning regulatory programs from functional genomics tracks (Linder et al., 2025). However, RNA abundance also reflects RNA stability, RBP binding, and processing, suggesting that Borzoi-style models could be tuned for RNA dynamics by incorporating direct measurements of post-transcriptional control.

We collected RNA-seq degradation time-course training tracks from human and mouse systems (Sasse et al., 2026) and paired these data with ecCLIP profiles, in addition to broader ENCODE eCLIP resources for half-life-associated RBPs (Van Nostrand et al., 2020) and Perturb-seq measurements for 3^′^-end processing regulators (Linder et al., 2025). We combined these with ENCODE DNase-seq/RNA-seq and FANTOM5 CAGE data for matching cell types (The ENCODE Project Consortium, 2012; Noguchi et al., 2017). We included the transcriptional readouts because steady-state RNA abundance is set jointly by synthesis and decay, and the two are confounded in expression data alone. A model that learns initiation rate from sequence can account for the transcriptional component of expression and attribute the remaining variation to post-transcriptional control, which sharpens the estimate of how much of RNA fate is set by stability. The resulting 40-million-parameter transformer U-net predicts RBP binding, RNA stability, and steady-state expression across 452 human and 160 mouse tracks from 393-kb input sequences and was trained as an ensemble of six replicate models (**Fig. 2d**). This multi-task design allows Borzuki to score sequence changes across linked post-transcriptional readouts, including expression, degradation, and RBP binding, rather than requiring separate specialized models for each output.

Borzuki accurately predicted held-out RNA-seq and CLIP-seq (i.e. both eCLIP and ecCLIP) coverage profiles. In ablations evaluated against HEK293T 4sU-based half-life measurements, adding ecCLIP profiles to the training data improved the prediction of RNA-seq degradation time-course tracks, and adding ENCODE eCLIP and 3^′^-regulator Perturb-seq measurements further improved performance (**Fig. 2e**). For the held-out gene ZNF611 in a K562 actinomycin D transcriptional shutoff experiment, in which inhibition of transcription allows RNA loss to be followed over time, predicted coverage recapitulated both gene-level log-coverage and position-specific signal across time points (**Fig. 2f**; Melé et al., 2017). For both ecCLIP and ENCODE eCLIP tracks, predicted and measured coverage agreed at gene-level and within-gene bin-level resolution (**Fig. 2g**; **Supp. Fig. S2c**), with ELAVL1 examples shown in **Supplementary Fig. S2e,f**. These analyses establish that the model can learn post-transcriptional regulatory tracks from sequence across held-out loci.

In a complementary comparison across model sizes and training-data compositions, models trained with CLIP-seq and Perturb-seq data consistently outperformed matched models trained without these modalities when predicting RNA-seq coverage, with the strongest performance observed for the 40-million-parameter model trained on the full dataset (**Fig. 2h**). Using K562 actinomycin D half-life tracks, Borzuki-predicted mRNA half-lives correlated with experimental measurements (Spearman R = 0.745) and with Saluki’s cell type-agnostic half-life predictions (Spearman R = 0.728; **Supp. Fig. S2d**; Agarwal and Kelley, 2022).

To extend the data-utility assessment to regulatory variation, we asked whether the learned features generalize to sequence perturbations not used for model training. Across fine-mapped 3^′^ UTR eQTLs, Borzuki distinguished high-confidence causal variants from matched negatives more effectively than Saluki (AUPRC = 0.694 for Borzuki vs 0.626 for Saluki), with the strongest performance from the 40-million-parameter model trained with the full CLIP- and Perturb-seq-expanded data set (**Fig. 2i**; Kerimov et al., 2021; Lonsdale et al., 2013). Borzuki also performed well on causal Mendelian 3^′^ UTR variants from TraitGym, particularly when variants were scored using the gene context with the largest predicted effect rather than only the annotated target gene (AUPRC = 0.545 for Borzuki vs 0.422 for Saluki) (**Supp. Fig. S2g**; Benegas et al., 2025). We further assessed variant scoring capabilities using massively parallel reporter assay (MPRA) data measuring 3^′^ UTR variant effects on RNA stability (**Supp. Fig. S2h**; Siegel et al., 2022). To score variants, we inserted reference and variant oligos in native 3^′^ UTRs and predicted differences in exon-aggregated log-coverage across RNA-seq and CLIP-seq tracks using the Borzuki ensemble (Methods). Borzuki was less accurate than Saluki on this task, but averaging the two models’ scores outperformed either model alone, suggesting that they capture complementary aspects of variant effects (**Supp. Fig. S2i**). Importantly, model ablations trained without ecCLIP data performed worse at half-life variant effect prediction (**Supp. Fig. S2j**). The ablation directly supports the contribution of ecCLIP to RNA-stability variant prediction; the eQTL and Mendelian comparisons describe the performance of the combined model on those specific benchmarks.

Together, these analyses establish ecCLIP as a high-quality, high-resolution resource for RBP-RNA interaction mapping. Our dataset shows concentrated signal at inferred binding sites, recovers independently measured RBP sequence preferences, and provides information that improves sequence-to-function modeling of RNA dynamics. The ablations demonstrate that the knowledge of RBP binding sites provided by ecCLIP adds predictive information about RNA half-life and RNA-stability variant effects.

### 2.3. Modeling RBP recognition and regulatory variation with Pythia

Having established the quality and utility of ecCLIP, we next used its nucleotide-resolution maps to learn the sequence and structural grammar that underlie RBP-RNA interactions. This problem is more complex than DNA motif discovery because RBP binding is shaped not only by primary RNA sequence, but also by basepairing, accessibility, and local folding context. Linear motif models, including k-mers, regular expressions, and position weight matrices, provide a natural representation for sequence preferences. In contrast, RNA structural elements couple distant nucleotide positions, making context-free grammars (CFGs) a natural formalism for representing paired sequence constraints. This idea has a long history in RNA modeling, including stochastic CFG approaches and RNApromo for discovering structural RNA motifs (Searls, 2002; Rabani et al., 2008), and was central to our earlier work where structural cis-regulatory elements were represented as families of CFGs and scored for association with post-transcriptional regulatory programs (Goodarzi et al., 2014, 2012; Fish et al., 2021).

We developed Pythia to bring this CFG-inspired representation into a neural network architecture for RNA-protein interaction modeling. Pythia uses two parallel arms: a sequence arm that processes one-hot encoded RNA sequence with trainable convolutional layers, and a structure arm that first passes the same sequence through a fixed dilated convolutional layer (**Fig. 3a**). The fixed layer enumerates candidate Watson-Crick and wobble base-pairing rules across a range of distances and bulge configurations. Conceptually, each kernel corresponds to a CFG production rule: a paired nucleotide constraint at a defined separation, with intervening positions left unconstrained. Because these kernels are frozen rather than learned, they provide an explicit structural inductive bias without requiring an external folding algorithm, experimentally measured structure, or task-specific structure labels. Downstream trainable layers then learn which combinations of these candidate structural rules are informative for a given supervised task.

**Figure 3.**
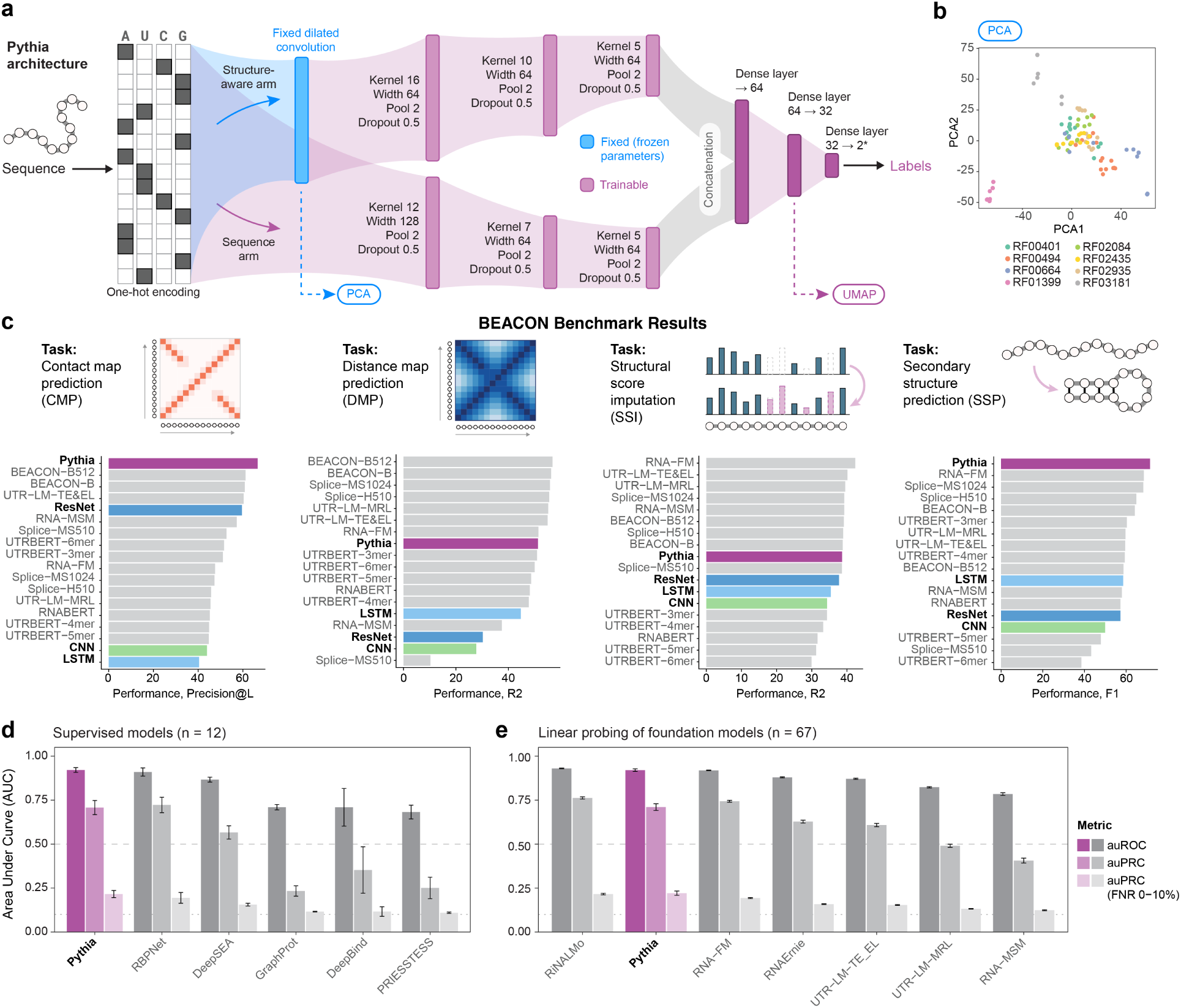
Pythia leverages the context-free grammar of RNA secondary structure. **(a)** Model architecture. Pythia incorporates a fixed dilated convolutional layer with frozen parameters (blue) that captures hierarchical structural patterns in RNA. The resulting representation is processed alongside raw sequence input by additional convolutional layers and three downstream dense layers for task-specific prediction. **(b)** RNA structural family organization. The first principal components of Pythia’s fixed convolutional layer embeddings cluster members of canonical RFAM RNA structural motif families, reflecting inherent structural grammar capture. **(c)** Benchmarking results across four BEACON RNA structure tasks; Contact Map Prediction (CMP), Distance Map Prediction (DMP), Structural Score imputation (SSI), and Secondary Structure prediction (SSP). Pythia achieves the top performance in CMP and SSP, while remaining highly competitive in DMP and SSI. **(d)** Binding-site prediction compared with five supervised baselines (RBPNet, DeepSEA, DeepBind, PRIESSTESS, and GraphProt) across the 12 RBPs shown. **(e)** Binding-site prediction compared with supervised linear probes on six frozen RNA foundation models (RiNALMo, RNA-FM, UTR-LM-TE_EL, UTR-LM-MRL, RNA-MSM, and RNAErnie; n=67 RBPs). For the binding tasks, supervised baselines and linear probes use the same labeled data splits as Pythia; foundation-model pretraining exposure is not controlled. For each model, bars from most opaque to most transparent show auROC, auPRC, and auPRC in the high-recall region (false-negative rate ≤ 10%), respectively. Models are ordered by mean auROC.

We first evaluated whether this representation accurately captured RNA structural information independently of RBP binding. When sequences from canonical RFAM structural families were passed through the fixed dilated convolutional layer, principal component analysis separated families with distinct structural organization (**Fig. 3b**; Kalvari et al., 2021). We also evaluated synthetic sequences containing an injected CFG-defined structural element and matched controls lacking the element, without fixed primary sequence identity. Their embedding distributions overlapped in the UMAP projection (**Supp. Fig. S3a**), which alone does not establish classification performance. As a more challenging structural classification task, we tested whether Pythia could distinguish preserved from disrupted pseudoknots generated by in silico mutation. Pythia achieved high performance on this task (AUROC = 0.95, AUPRC = 0.87), outperforming supervised sequence models including LSTM, DeepBind, DeepSEA, and ResNet architectures (**Supp. Fig. S3b,c**; Alipanahi et al., 2015; Zhou and Troyanskaya, 2015; He et al., 2016).

To benchmark the architecture more broadly, we evaluated Pythia on four RNA structural tasks of the BEACON suite: contact map prediction (CMP), distance map prediction (DMP), structural score imputation (SSI), and secondary structure prediction (SSP; Ren et al., 2024). The 17 BEACON baselines comprise conventional CNN, ResNet, and LSTM architectures; pretrained RNA language models, including RNA-FM, RNABERT, RNA-MSM, SpliceBERT, 3UTRBERT, and UTR-LM variants; and BEACON-B/BEACON-B512 (Methods). Against this panel, Pythia achieved the top overall performance on the two contact and secondary-structure prediction tasks: CMP (Top-L precision 66.6%) and SSP (F1 71.5%), surpassing even the purpose-built BEACON-B baseline. For SSI (R^2^ 38.6%) and DMP (R^2^ 51.3%), Pythia exceeded the supervised CNN, ResNet, and LSTM baselines, while trailing the top pretrained RNA language models on DMP (**Fig. 3c**). Together, these benchmarks show that Pythia’s fixed base-pairing prior captures RNA secondary-structure constraints, and motivated our application of Pythia to RBP binding sites from ecCLIP data to simultaneously capture the sequence and structural determinants of RBP-RNA interactions.

### 2.4. Pythia predicts ecCLIP binding sites from sequence and structural context

We next trained Pythia on the ecCLIP atlas to predict RBP binding sites. For each of the 67 RBPs, we trained a separate binary classifier to discriminate ecCLIP peaks from dinucleotide-shuffled versions of the same sequences, which serve as composition-matched negatives, with held-out chromosomes used for evaluation. Thus, RBP identity was not provided as a model input; instead, each model learned an RBP-specific binding function using the same sequence-plus-structure architecture. This design allowed us to test whether the fixed structural arm improved binding-site prediction while preserving a common framework for comparing sequence and structure contributions across RBPs. Per-RBP auROC and auPR values summarize variation in prediction performance across the atlas (**Supp. Fig. S3e**).

We compared Pythia with five supervised baselines (DeepBind, DeepSEA, GraphProt, PRIESSTESS, and RBPNet) and with supervised linear probes on frozen embeddings from six RNA language models (RiNALMo, RNA-FM, UTR-LM-TE_EL, UTR-LM-MRL, RNA-MSM, and RNAErnie). Across matched RBP binding tasks, Pythia outperformed the five baselines trained on sequence features and/or externally supplied structure features in mean auROC (0.92 vs. 0.91 for the closest, RBPNet; **Fig. 3d**; Maticzka et al., 2014; Laverty et al., 2022; Her et al., 2026). For EIF2AK2, an example RBP with clear separation among models, Pythia achieved auROC = 0.89 and auPR = 0.61, compared with auROC = 0.85 and auPR = 0.60 for the next-best supervised comparator, RBPNet (**Supp. Fig. S3f**). This advantage was largest when training data were limited: with 10% of the training data, Pythia achieved a mean auROC of 0.82, compared with 0.74 for RBPNet and 0.56–0.73 for the other baselines, outperforming every baseline on every RBP evaluated, and retained the highest precision in the high-recall regime (**Supp. Fig. S3g**). For the embedding comparisons, Pythia and the linear probes used matched labeled data splits, but the foundation models may have encountered test sequences during pretraining. Thus, labeled-data supervision is matched while pretraining exposure is not controlled. With that caveat, Pythia performed comparably to the strongest foundation-model embeddings, ranking second by mean auROC after RiNALMo, while retaining a compact supervised architecture with an explicit structural representation (**Fig. 3e**). These results show that Pythia learns accurate RBP binding-site predictors from the ecCLIP atlas and that an explicit structural inductive bias can match or exceed learned sequence representations on this task. Pythia’s downstream weights are learned with supervision and the fixed component is the initial base-pairing feature layer.

### 2.5. Pythia attribution and MPRC test structural determinants of RBP binding

Having trained Pythia to predict ecCLIP binding sites, we next asked whether the model could identify which binding events depend primarily on RNA sequence and which depend on structural context. Because Pythia contains separate sequence and structure arms, we can quantify the contribution of each arm to individual RBP binding predictions. For each RBP-specific model, we applied DeepLIFT separately to the sequence arm and the fixed-dilated-convolution structure arm, comparing the effect of perturbing each input representation on the predicted binding score (Shrikumar et al., 2019). The resulting sequence-to-structure attribution ratio summarized the model’s relative reliance on each arm at each binding site, with high ratios corresponding to sequence-dominant predictions and low ratios corresponding to structure-dominant predictions (**Fig. 4a**).

**Figure 4.**
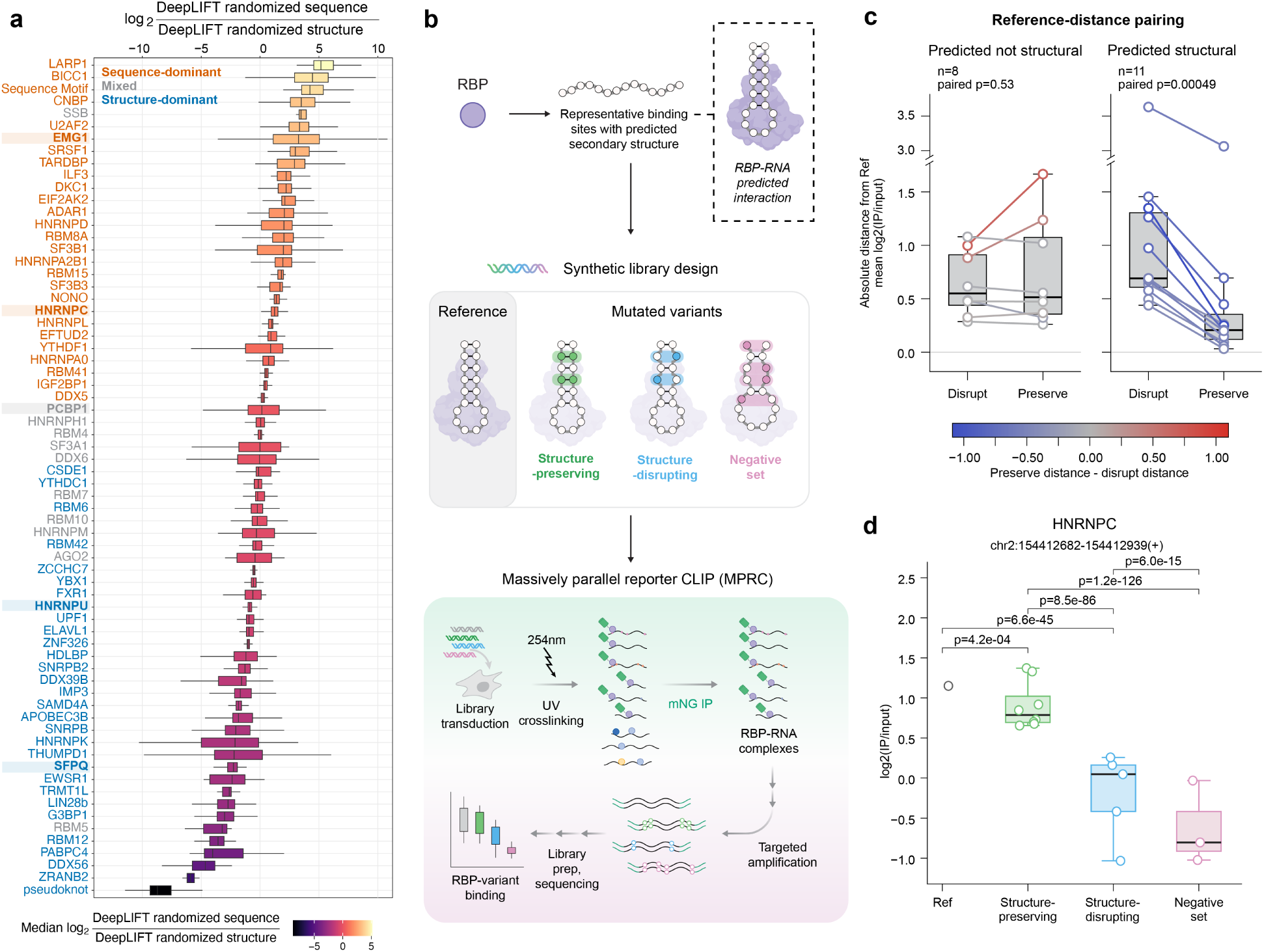
DeepLIFT and massively parallel reporter CLIP validate structure-dependent RBP binding determinants. **(a)** DeepLIFT-based sequence and structure attribution across RBP binding models. For each RBP-specific Pythia model, DeepLIFT scores were computed after perturbing the sequence arm or the fixed-dilated-convolution structure arm. The log_2_ sequence-to-structure attribution ratio summarizes relative model reliance on each arm. Each box shows, for one RBP, the distribution of per-peak log_2_ ratios across its ecCLIP peaks, restricted to the 10% of peaks with the highest variance across the sequence, structure, and full-model DeepLIFT scores (Methods). Boxes show the median and interquartile range, whiskers extend to 1.5x the interquartile range, outliers are not shown, and fill color indicates the RBP’s median log_2_ ratio. RBP-level labels are derived from these Pythia scores: an absolute median ratio >0.05 and Benjamini-Hochberg FDR <10% for a one-sample Wilcoxon test against zero define sequence-dominant (positive) or structure-dominant (negative) models; the remaining models are labeled mixed (Methods). Fixed sequence motifs and pseudoknot-derived controls illustrate the positive and negative directions of the attribution scale, respectively. **(b)** Schematic of massively parallel reporter CLIP (MPRC). Pythia-nominated RBP binding sites are synthesized as pooled reporter libraries containing reference sequences, structure-preserving mutations, structure-disrupting mutations, and multi-mutation negative controls. Reporter libraries are introduced into HEK293T cells expressing the corresponding endogenously tagged RBP, and reporter-targeted CLIP is used to quantify relative RBP binding. **(c)** Paired reference-distance analysis of MPRC validation sites whose reference reporters retained binding. Of 30 structure-validation sites tested, 19 showed reference enrichment (Ref log_2_(IP/input) > 0.25) and were included in this analysis. For each reference-enriched site, the absolute distance from the reference log_2_(IP/input) was compared for structure-disrupting and structure-preserving reporters. Sites predicted by Pythia to be structure-dependent showed significantly smaller reference distance for structure-preserving reporters than structure-disrupting reporters (n = 11, one-sided paired Wilcoxon P = 4.9 × 10^−4^), whereas sites not predicted to be structure-dependent showed no significant paired difference (n = 8, P = 0.527). **(d)** Representative HNRNPC validation site. For an intronic GALNT13 site at chr2:154,412,682-154,412,939(+), structure-preserving mutations maintain binding near the reference sequence, whereas structure-disrupting mutations and multi-mutation negative controls show reduced binding.

Across the ecCLIP atlas, DeepLIFT revealed substantial heterogeneity in the sequence and structure contributions to RBP binding. Positive-control sequences with fixed primary-sequence motifs showed strong sequence dominance, whereas pseudoknot-derived structural controls showed strong structure dominance, providing controls for the direction of the attribution scale (**Fig. 4a**). We assigned RBP-level sequence-dominant or structure-dominant labels from Pythia attributions, requiring an absolute median log_2_ sequence-to-structure ratio > 0.05 and a one-sample Wilcoxon test against zero with Benjamini–Hochberg FDR < 10% (Methods). Models that did not meet these criteria were labeled mixed; the site-level scores nevertheless form a continuum: some RBPs showed predominantly sequence-driven binding, others showed stronger structural dependence, and many exhibited site-specific variation. This heterogeneity indicates that structural dependence is not only an RBP-level property: individual sites bound by the same RBP can differ in the degree to which Pythia relies on primary sequence versus structural context.

To test whether these model-derived attributions correspond to experimentally measurable binding determinants, we developed massively parallel reporter CLIP (MPRC). For five RBPs spanning the sequence-structure continuum (EMG1, HNRNPC, PCBP1, HNRNPU, and SFPQ), we selected representative ecCLIP binding sites and designed synthetic reporter libraries containing the reference sequence together with targeted mutant alleles (**Fig. 4b,c**). For each site, the library included mutations designed to preserve the predicted RNA structure, mutations designed to disrupt the predicted structure, and multi-mutation negative controls. We introduced these libraries into HEK293T cells expressing the corresponding endogenously tagged RBP and measured relative RBP binding by reporter-targeted CLIP. This design allowed us to directly test whether sites predicted to depend on structural context were specifically sensitive to structure-disrupting mutations while retaining binding under structure-preserving mutations.

We quantified the MPRC output using UMI-deduplicated reporter counts, normalized IP and input libraries, and summarized binding for each construct as log_2_(IP/input). Because these reporters move endogenous binding sites outside their native transcript context, we first asked whether the reference reporter retained measurable binding in the CLIP assay. Of 30 structure-validation sites tested, 19 showed clear reference enrichment (Ref log_2_(IP/input) > 0.25), including 11 of 18 sites predicted to be structure-dependent and 8 of 12 sites not predicted to be structure-dependent. We restricted the rescue analysis to this reference-enriched subset and grouped reporters as reference, structure-preserving, structure-disrupting, or negative-control constructs. Read-depth distributions for the reference-enriched sites were comparable between predicted structure-dependent and not-structure-dependent classes, and rescue effects were not restricted to low-count sites (**Supp. Fig. S4a**). Because a nonsignificant difference from the reference can reflect limited power rather than true preservation of binding, we used effect-size rescue as the primary validation statistic. Specifically, we asked whether structure-preserving mutations remained closer to the reference log_2_(IP/input) than structure-disrupting mutations.

MPRC confirmed that Pythia-nominated structural features correspond to functional determinants of RBP binding. Among sites predicted by Pythia to be structure-dependent, structure-preserving reporters were closer to the reference than structure-disrupting reporters at all 11 reference-enriched sites, using the same number of mutations in both conditions (one-sided paired Wilcoxon P = 4.9 × 10^−4^; **Fig. 4c**). In contrast, sites not predicted to be structure-dependent showed no significant paired difference (n = 8, P = 0.527; **Fig. 4c**).

We additionally tested whether structure-preserving and structure-disrupting constructs differed in IP/input enrichment using binomial logistic regression adjusted for replicate (Methods). Requiring a significant difference between these groups yielded the expected pattern at 10 of 10 predicted structure-dependent sites with valid model fits, compared with 4 of 8 sites not predicted to be structure-dependent (one-sided Fisher exact P = 0.023). At the continuous-score level, more structure-dependent Pythia scores were associated with stronger rescue by structure-preserving mutations (**Supp. Fig. S4b**), and additional representative validation sites across HNRNPC, PCBP1, and EMG1 illustrated the range of responses across sites (**Supp. Fig. S4c– f**). One nominated example was an intronic GALNT13 sequence bound by HNRNPC, in which the reference sequence and structure-preserving variants showed strong enrichment (1.15 and 0.90 mean log_2_(IP/input), respectively), whereas structure-disrupting and negative-control variants showed reduced binding (−0.20 and −0.62 mean log_2_(IP/input); **Fig. 4d**). This example is consistent with previous work showing that local RNA structural changes can expose HNRNPC binding motifs and modulate HNRNPC-RNA interactions (Liu et al., 2015). Predicted RNA structures illustrate possible changes in poly-U accessibility in this reporter, providing structural context for the binding measurements (**Supp. Fig. S4g**).

These results support the use of Pythia’s structure-arm attributions to nominate binding sites that are sensitive to structure-disrupting mutations in the reporter context. More broadly, they establish that structure-aware interaction models can nominate specific RNA structural features whose disruption alters binding while compensatory or structure-preserving sequence changes retain activity.

### 2.6. Pythia models predict allele-specific effects of trait-associated variants on RBP binding

Having shown that RBP-specific Pythia models can nominate structural determinants of RBP binding, we next asked whether the same model collection could predict the effects of naturally occurring genetic variants on RBP binding. This is a central use case for structure-aware RBP binding models: many disease- and trait-associated variants fall in transcribed noncoding regions, but their effects on post-transcriptional regulation are difficult to interpret from sequence context alone. We therefore focused on phenotype-annotated variants located within ecCLIP binding sites for five RBPs assayed in the MPRC validation framework (EMG1, HNRNPC, HNRNPU, PCBP1, and SFPQ). For each RBP-variant pair, we used the corresponding RBP-specific Pythia model to score the reference and alternate alleles and prioritized variants spanning predicted losses, gains, and minimal effects on binding (**Fig. 5a,b**).

**Figure 5.**
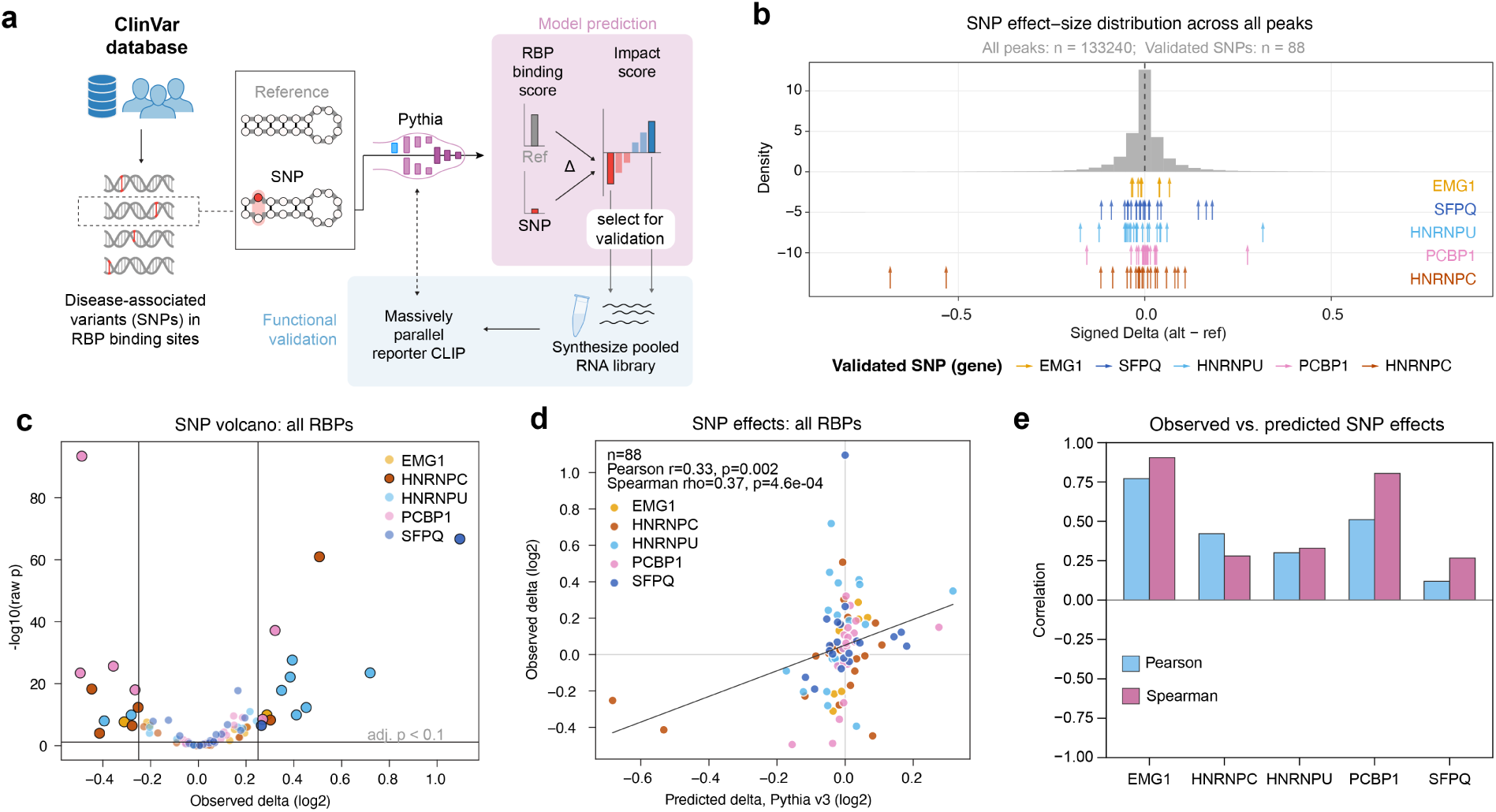
RBP-specific Pythia models predict allele-specific effects of trait-associated variants on RBP binding. **(a)** Schematic of the targeted variant-validation workflow with MPRC. Phenotype- or trait-annotated variants overlapping ecCLIP binding sites are scored with RBP-specific Pythia models, reference and alternate alleles are cloned into pooled reporter libraries, and MPRC is used to quantify allele-specific RBP binding. **(b)** Variant selection strategy. For each RBP, the distribution of allele effects predicted by the corresponding Pythia model is shown for candidate variants overlapping ecCLIP binding sites, with selected predicted-loss, predicted-gain, and low-effect variants highlighted. **(c)** MPRC measurements of allele-specific binding. Volcano plot shows the observed allelic effect, defined as log_2_(IP/input)_Alt_ -log_2_(IP/input)_Ref_, versus -log_10_(raw P) from replicate-adjusted binomial logistic regression. Variants passing global FDR < 0.1 and |observed effect| > 0.25 are highlighted. **(d)** Concordance between predicted and experimentally measured variant effects across all tested RBP-variant pairs. Each point represents one RBP-variant pair, colored by RBP; predictions were generated by the RBP-specific Pythia model corresponding to the assayed RBP. Across 88 tested pairs, predicted and observed effects were positively correlated (Pearson r = 0.33, P = 0.0019; Spearman rho = 0.37, P = 4.6 × 10^−4^). **(e)** RBP-specific concordance between predicted and observed variant effects. Bars show Pearson and Spearman correlations for EMG1 (n = 8), HNRNPC (n = 20), HNRNPU (n = 20), PCBP1 (n = 20), and SFPQ (n = 20).

We experimentally tested 88 RBP-variant pairs using the same pooled reporter CLIP strategy (i.e. MPRC). Reference and alternate 50-nt sequences were synthesized into a pooled library, introduced into HEK293T cells expressing the corresponding endogenously tagged RBP, and quantified by MPRC. We then compared allele-specific RBP enrichment between each matched reference and alternate sequence testing whether the alternate allele changes RBP enrichment relative to its matched reference sequence while controlling for input abundance and replicate effects.

Through MPRC, we identified allele-specific RBP binding effects across all five RBPs. Using a global FDR threshold of 10% and an absolute effect-size threshold of 0.25 log_2_ units, 24 of 88 tested RBP-variant pairs showed significant allele-specific binding (**Fig. 5c**). Variant effects predicted by the RBP-specific Pythia models were positively correlated with experimentally measured effects across the full validation set (Pearson r = 0.33, P = 0.0019; Spearman *ρ* = 0.37, P = 4.6 × 10^−4^; **Fig. 5d**). Directional concordance was also above chance with 53 of 86 non-zero predicted variants showing concordant direction (one-sided binomial P = 0.020).

Performance varied across RBPs, with the strongest concordance for PCBP1 (Pearson r = 0.51, P = 0.021; Spearman *ρ* = 0.80, P = 1.9 × 10^−5^) and EMG1 (Pearson r = 0.77, P = 0.025; Spearman *ρ* = 0.90, P = 0.002), and weaker correlations for HNRNPC, HNRNPU, and SFPQ (**Fig. 5e**). This RBP-level heterogeneity reinforces the need to pair model-based variant prioritization with experimental calibration in the relevant assay system. These measurements evaluate Pythia’s variant predictions in the selected reporter set; the binding-site classification benchmarks in **Fig. 3** do not establish comparative variant-effect performance for the other models. Observed effects stratified by predicted direction and examples with large predicted effects are shown in **Supplementary Fig. S5a,b**.

To highlight a few examples, we applied a strict concordance filter requiring |predicted effect| > 0.1 log_2_ units, significant allele-specific binding, and matched predicted and observed direction (**Supp. Fig. S5b**). This highlighted a small set of functional candidates with trait-relevant annotations. Two intronic variants were predicted and validated to reduce HNRNPC binding: rs9323624, whose locus is annotated to non-steroidal anti-inflammatory drug-induced acute urticaria and angioedema, and rs56302494, annotated to heel bone mineral density. Because HNRNPC binding can influence RNA processing, these variants nominate local RNA-processing hypotheses for follow-up in native transcript and disease-relevant cellular contexts. The same strict filter also identified a gain of HNRNPU binding at the BCKDHB-locus variant rs9361614, annotated to waist circumference adjusted for body mass index, and loss of PCBP1 binding at rs11668313, annotated to low-density lipoprotein cholesterol levels. These links are speculative and should not be interpreted as causal disease mechanisms; rather, they illustrate how allele-specific RBP binding measurements can generate testable hypotheses for mechanistic dissection of trait-associated loci.

These results show that an ecCLIP-trained collection of RBP-specific, structure-aware models can prioritize naturally occurring variants that alter RBP binding in a pooled experimental assay. The validation should be interpreted specifically as evidence for allele-specific binding effects in the HEK293T cellular context, rather than as direct evidence of disease mechanism in the relevant tissue or cell type. Nonetheless, by linking trait-associated sequence variation to measured changes in RBP binding, this analysis establishes a practical route from ecCLIP-trained, structure-aware prediction to experimentally testable post-transcriptional regulatory hypotheses.

## 3. Discussion

RNA-binding proteins regulate gene expression through sequence recognition, RNA structural context, transcript position, and cellular state. Dissecting these mechanisms requires experimental maps that are uniform enough for atlas-scale comparison and resolved enough to support quantitative modeling. Here we developed eclair CLIP (ecCLIP), a scalable CLIP-seq framework that combines endogenous split-fluorescent-protein tagging with ligation-free library preparation. Applied to 67 RBPs in HEK293T cells, ecCLIP generated a high-quality RBP binding atlas that recovers known sequence preferences, captures regulatory signal linked to RNA half-life, and provides a substrate for structure-aware modeling of RBP-RNA interaction grammar.

A central technical feature of ecCLIP is that endogenous tagging creates a common and stringent affinity-capture strategy across RBPs while preserving endogenous expression control. Endogenous epitope tagging was previously used in TAG-eCLIP to enable profiling without RBP-specific antibodies (Van Nostrand et al., 2017). ecCLIP combines endogenous split-fluorescent-protein tagging with infrared visualization and ligation-free library preparation. This addresses two persistent limitations of atlas-scale CLIP studies: antibodies differ substantially in availability, specificity, and capture efficiency, whereas approaches based on exogenous tagged constructs can perturb RBP abundance, stoichiometry, localization, or binding behavior (Ule et al., 2003; König et al., 2010; Van Nostrand et al., 2020). The shared endogenous capture handle does not remove all sources of variation, but it reduces major RBP-specific technical variables without requiring ectopic over-expression from a transgene. Because many profiled RBPs were selected from the OpenCell collection, ecCLIP also adds a transcriptome-wide RNA-occupancy layer to a broader endogenous-tagging resource that includes protein localization and interaction measurements (Cho et al., 2022). This creates opportunities to interpret RBP binding alongside other OpenCell modalities while contributing a complementary data type to this resource.

The atlas shows that high-resolution RBP occupancy profiles are useful beyond binding-site annotation. Incorporating ecCLIP tracks into Borzoi-style models improved prediction of HEK293T RNA half-life, and expanding the training set with broader CLIP and Perturb-seq data improved it further. Borzuki serves here as a test of data utility: it extends the Borzoi framework to incorporate these measurements into prediction of RNA stability, RBP binding, and expression dynamics from genomic sequence (Linder et al., 2025). The ecCLIP ablations on RNA half-life and MPRA variant effects provide direct evidence for the added information in these binding maps. Performance on fine-mapped eQTLs and causal Mendelian 3^′^ UTR variants further assesses the combined model, while the MPRA results show complementarity with Saluki. These results should still be viewed as a step toward, rather than a complete solution for, post-transcriptional variant interpretation, because 3^′^ UTR variant effects remain dependent on cell type, assay, molecular readout, and transcript context.

Pythia provides the modeling framework for asking which sequence and structural features determine RBP recognition and how their perturbation changes binding. Deep learning models of transcription factor-DNA binding are motivated not only by prediction in new sequences, but also by the possibility of learning regulatory syntax from functional genomics data (Alipanahi et al., 2015; Kelley et al., 2018; Avsec et al., 2021). RBP-RNA interactions add a structural layer because binding can depend on both primary sequence and RNA accessibility or pairing. Building on prior work showing that structural context shapes post-transcriptional regulatory elements, including TEISER and pyTEISER (Goodarzi et al., 2012), Pythia explicitly separates sequence and structure arms so that model predictions can be decomposed into these components. The DeepLIFT and MPRC analyses show that this decomposition is experimentally meaningful: structure-preserving mutations were more likely to retain binding than structure-disrupting mutations at Pythia-nominated sites. The HNRNPC/GALNT13 example illustrates this logic, with structure-preserving mutations retaining HNRNPC binding and structure-disrupting mutations reducing binding, consistent with prior evidence that local RNA structure modulates HNRNPC accessibility (Liu et al., 2015).

RBP-specific Pythia models also prioritized naturally occurring variants with allele-specific binding effects in the reporter assay. Across 88 tested RBP-variant pairs, predicted and observed effects were positively correlated, and directional concordance was above chance after excluding variants with zero predicted effect. The strongest examples included HNRNPC binding loss at rs9323624 and rs56302494, HNRNPU binding gain at the BCKDHB-locus variant rs9361614, and PCBP1 binding loss at rs11668313. These findings should be interpreted as molecular effects on RBP binding in the HEK293T reporter system, not as causal disease mechanisms. Their trait annotations nominate hypotheses for follow-up, but establishing organism-level consequences will require testing native transcripts, relevant cellular states, and downstream RNA-processing or expression effects.

Several limitations define the next steps. The atlas was generated in HEK293T cells, so extending ecCLIP across cell types, cell perturbations, and disease contexts will be important for modeling context-specific regulation. Endogenous tagging provides a uniform capture strategy but still modifies each RBP locus, making orthogonal functional checks important for proteins sensitive to terminal tags. Pythia’s structure arm encodes structural potential rather than directly measured RNA structure, and reporter assays simplify native transcript context. Pairing ecCLIP with structure probing, long-read isoform measurements, perturbations of trans-acting RBPs, and endogenous genome editing will help connect local interaction grammar to native regulatory outcomes.

Together, these results establish ecCLIP as both an experimental atlas and a modeling substrate for post-transcriptional regulation. The data improve genome-scale modeling of RNA output through Borzuki and enable structure-aware interaction modeling through Pythia. As RNA-focused sequence-to-function models continue to expand, their utility will depend on the quality, resolution, and consistency of the biological measurements used to train and evaluate them. ecCLIP addresses this need for RBP-RNA interactions and provides a foundation for decoding how RNA sequence and structure jointly shape post-transcriptional gene regulation.

## 4. Methods

### 4.1. Generation of stable mNG-tagged RBP HEK293T cell lines

The 67 HEK293T cell lines expressing split mNeonGreen-tagged proteins were generated as previously described (Cho et al., 2022). The Methods Table lists the 67 proteins in the ecCLIP atlas; UniProt identifiers refer to the protein sequences used for analysis.

### 4.2. Cell Culture

HEK293T cell lines were cultured in Dulbecco’s Modified Eagle Medium supplemented with 10% FBS, 1*μ*g/mL Amphotericin (Corning), and 100 U/mL penicillin, 100 *μ*g/mL streptomycin (Corning) in a 5% CO_2_ humidified incubator.

### 4.3. eclair CLIP-seq

#### 4.3.1 Cell Crosslinking

Cells were harvested at 80-90% confluency, two 15cm plates were used per replicate. Cells were washed 1X with ice-cold PBS and then irradiated with 400mJ/cm^2^ 254nm UV. On ice, cells were collected by scraping, and cell pellets frozen at −80^◦^C.

#### 4.3.2 Immunoprecipitation

Cell pellets were lysed on ice in 600*μ*L cold low salt wash buffer (1X PBS, no Mg++ or Ca++, 0.1% SDS, 0.5% sodium deoxycholate, 0.5% IGEPAL CA-630) with 1X halt EDTA-free protease inhibitor (Thermo) and 6*μ*L of SuperaseIn RNase inhibitor (ThermoFisher). 60*μ*L DNase I (1 U/*μ*L, Promega) was added to the lysate and incubated at 37^◦^C for 10 minutes with shaking at 1000rpm. Lysate was then divided equally into 2 tubes. To tube 1, 10*μ*L of a high concentration RNase mix (3.3ng/*μ*L RNase A, 1 U/*μ*L RNase I) was added; to tube 2 10*μ*L of a low RNase mix (0.67ng/*μ*L RNase A, 0.2 U/*μ*L RNase I) was added. RNase mixes were diluted in low salt wash buffer before adding to lysate. Lysate was then incubated at 37^◦^C for 5 minutes. The two RNase-treated tubes were then combined and spun at 4^◦^C at 20,000xg for 20 minutes. The resulting clarified lysate was added to 25*μ*L anti-neon green magnetic beads (Proteintech) that had been pre-washed 2X with PBST (PBS + 0.02% tween-20) and 1X with low salt wash buffer. The bead-lysate mixture was rotated end-over-end for 1.5-2 hours at 4^◦^C. Beads were washed 2X with cold low salt wash buffer, 2X with cold high salt wash buffer (5X PBS no Mg++ or Ca++, 0.1% SDS, 0.5% sodium deoxycholate, 0.5% IGEPAL CA-630), and 2X with cold PNK wash buffer (50mM Tris-HCl pH 7.5, 10mM MgCl_2_, 0.5% IGEPAL CA-630).

#### 4.3.3 Dephosphorylation

After removing the final wash, RNA was dephosphorylated by resuspending the beads in 2.5*μ*L 10X PNK buffer (500mM Tris pH 6.8, 5mM MgCl_2_, 50mM DTT), 2*μ*L T4 PNK (10u/*μ*L, NEB), 0.5*μ*L SuperaseIn (Ther-moFisher), 20*μ*L H_2_O. Samples were incubated at 37^◦^C for 20 minutes with interval shaking at 1350rpm 15 seconds / 5 minutes off. Beads were washed 1X with cold PNK wash buffer, 1X with cold high salt wash buffer and 2X with PNK wash buffer.

#### 4.3.4 PolyA tailing RNP complexes

Following dephosphorylation, RNA was poly(A) tailed. On ice, beads were resuspended in 0.8*μ*L of yeast poly(A) polymerase (Jena 600U/*μ*L), 4*μ*L 5X yeast PAP buffer, 1*μ*L 10mM ATP (unlabeled), 0.5*μ*L SuperaseIn, and 13.7*μ*L H_2_O. Samples were then incubated at 22^◦^C for 5 minutes with a 15 second shake at 1350rpm after 2.5 minutes. The beads were then washed 2X with cold high salt wash buffer and 2X with cold PNK wash buffer.

#### 4.3.5 Azide end labeling of RNP complexes

RNA was end labeled with azide by adding a mix containing 0.4*μ*L yeast poly(A) polymerase (Jena 600U/*μ*L), 2*μ*L 5X yeast PAP buffer, 0.25*μ*L SuperaseIn, 2*μ*L 10mM 2^′^-Azido-2^′^-Deoxyuridine-5^′^-Triphosphate (TriLink BioChemicals), and 5.35*μ*L H_2_O to the beads. Samples were incubated at 37^◦^C for 20 minutes and interval shaking at 1350rpm 15 seconds / 5 minutes off. Beads were washed 2X with cold high salt wash buffer and 2X with cold 1X PBS

#### 4.3.6 IR labeling of RNP complexes

Samples were then infrared labeled by adding 20*μ*L of 0.2mM 800CW DBCO (LiCor Biosciences) in PBS. Beads were incubated at 22^◦^C for 30 minutes with interval shaking at 1350rpm for 15 seconds / 5 minutes off, protected from light. Beads were then washed 1X with cold high salt wash buffer and 1X with cold PNK wash buffer. After removal of final wash, beads were resuspended in loading buffer (1X NuPAGE loading buffer (ThermoFisher), 50mM DTT). Beads were incubated at 75^◦^C for 10 minutes shaking at 1000rpm protected from light. Beads were magnetized and the eluate collected.

#### 4.3.7 PAGE Gel and Transfer

The eluates were run on a NuPAGE 4-12% Bis-Tris gel in MOPS running buffer (ThermoFisher) at 4^◦^C, protected from light. Samples were transferred to Protran BA-85 nitrocellulose membrane (Whatman) in 1X NuPAGE transfer buffer (ThermoFisher) with 10% ethanol. Membrane was imaged on a LI-COR Odyssey instrument. To isolate the RNA bound to the protein of interest, the region ∼20-45kDa above the molecular weight of the protein of interest was excised from the membrane.

#### 4.3.8 Proteinase K digest and RNA capture

RNA was released by submerging the membrane pieces in 200*μ*L proteinase K digestion buffer (100mM Tris-HCL pH 7.5, 100mM NaCl, 1mM EDTA, 0.2% SDS), and 12.5*μ*L proteinase K (Ambion, 20U/*μ*L) and incubating at 55^◦^C for 45 minutes with shaking at 1100rpm. The eluate was transferred to clean tubes and the NaCl concentration of the eluate was adjusted to 0.5M NaCl. This eluate was then transferred to oligo d(T) magnetic beads (NEB) that had been pre-washed 2X with proteinase K buffer. Samples were then incubated at 25^◦^C for 20 minutes with interval shaking at 300rpm 10 min / 1350rpm 15 seconds. Beads were then washed 2X with cold high salt wash buffer and 2X with cold 1X PBS. Samples were eluted off of the beads by adding 8*μ*L of bead elution buffer (20mM Tris-HCl pH 7.5, 1mM EDTA) and incubating at 50^◦^C for 5 minutes. The eluate was then transferred to PCR tubes.

#### 4.3.9 Library preparation

Library preparation was performed using the Takara SMARTer smRNA-Seq Kit with modifications. To the RNA from the previous step, 2.5*μ*L smRNA Mix 1 and 1*μ*L of 10*μ*M ecCLIP UMI RT primer was added and incubated at 72^◦^C for 3 minutes followed by 2-5 minutes on ice. Then 6.5*μ*L kit smRNA Mix 2, 0.5*μ*L RNase inhibitor, and 2*μ*L PrimeScript RT (200U/*μ*L) was added and samples incubated at 42^◦^C for 60 minutes, 70^◦^C for 10 minutes, and then placed back on ice. To this cDNA, 50*μ*L of 2X SeqAmp CB PCR Buffer, 2*μ*L SeqAmp DNA Polymerase, 2*μ*L 10*μ*M ecCLIP LP reverse primer and 24*μ*L H_2_O was added. Samples were indexed by adding 2*μ*L of a unique 10*μ*M Takara forward index primer (provided in kit) to each sample. Samples were amplified using the following program: 98^◦^C 1 minute followed by 98^◦^C 10 seconds -60^◦^C 5 seconds -68^◦^C 10 seconds, repeated for the number of cycles determined by qPCR to ensure no overamplification. Libraries were cleaned up using SPRI beads at a 1.1X bead : PCR reaction ratio. Library size distribution was assessed with a Tapestation instrument (Agilent), and pooled libraries were sequenced on a HiSeq 4000 (Illumina) at the UCSF Center for Advanced Technologies.

### 4.4. Massively Parallel Reporter CLIP-seq (MPRC)

#### 4.4.1 CLIP reporter cell line construction

Sequences containing the binding site for the RBP of interest, along with mutant variants (Methods Table), were cloned immediately 3^′^ of an EGFP CDS in a lentiviral backbone. Using lentiviral transduction, each variant sequence library was stably integrated into a HEK293T cell line expressing the split mNeonGreen-tagged target RBP.

#### 4.4.2 Cell Crosslinking

Cells were harvested at 80-90% confluency, two 15cm plates were used per replicate. Cells were washed 1X with ice-cold PBS and then irradiated with 400mJ/cm^2^ 254nm UV. On ice, cells were collected by scraping, and cell pellets frozen at −80^◦^C.

#### 4.4.3 Immunoprecipitation

Cell pellets were lysed on ice in 1.2mL cold low salt wash buffer (1X PBS, no Mg++ or Ca++, 0.1% SDS, 0.5% sodium deoxycholate, 0.5% IGEPAL CA-630) with 1X halt EDTA-free protease inhibitor (Thermo) and 12*μ*L of SuperaseIn RNase inhibitor (ThermoFisher). 120*μ*L DNase I (1 U/*μ*L, Promega) was added to the lysate and incubated at 37^◦^C for 10 minutes with shaking at 1000rpm. The lysate was spun at 4^◦^C 20,000xg for 20 minutes. 5% of the resulting clarified lysate was retained to use as the input sample. The remaining clarified lysate was added to 25*μ*L anti-neon green magnetic beads (Proteintech) that had been pre-washed 2X with low salt wash buffer. The bead-lysate mixture was rotated end-over-end for 1.5 hours at 4^◦^C. Beads were then washed 3X with cold low salt wash buffer and then 3X with cold high salt wash buffer (5X PBS no Mg++ or Ca++, 0.1% SDS, 0.5% sodium deoxycholate, 0.5% IGEPAL CA-630).

#### 4.4.4 Proteinase K digest and RNA isolation

RNA was released from the beads by adding 200*μ*L proteinase K digestion buffer (100mM Tris-HCL pH 7.5, 100mM NaCl, 1mM EDTA, 0.2% SDS), and 12.5*μ*L proteinase K at 37^◦^C for 1 hour with interval shaking at 1100rpm 30 seconds / 5 minutes off. The lysate input samples were processed in parallel. Acid phenol chloroform (Sigma) extraction was used to isolate the RNA. The RNA was ethanol precipitated, washed 2X with 75% ethanol, air dried and resuspended in 18*μ*L H_2_O. The input RNA was resuspended in 54*μ*l H_2_O.

#### 4.4.5 Library preparation

cDNA was synthesized by mixing 1*μ*L of 2*μ*M reporter MPRC UMI RT primer, 1*μ*L 10mM dNTPs, and 11.5*μ*L of the IP RNA isolated above. For the input samples, 1*μ*g of RNA was used as input. Samples were incubated at 65^◦^C 5min then placed immediately on ice. To this, 4*μ*L 5X RT buffer, 1*μ*L 100mM DTT, 0.5*μ*L SuperaseIN and 1*μ*L SuperScript IV RT (ThermoFisher) were added. Samples were incubated at 55^◦^C 10min then 80^◦^C for 10 min. In parallel, reactions without reverse transcriptase were performed to assess for DNA contamination.

To digest free oligo, 1*μ*L thermolabile exonuclease I (20 U/*μ*L, NEB) was added to each sample and incubated at 37^◦^C 4 min then 80^◦^C 1 min. Libraries were amplified by combining 50*μ*L 2X KAPA HiFI HotStart Ready mix (Roche), 2*μ*L 10*μ*M reporter MPRC forward PCR1 primer, 2*μ*L 10*μ*M reverse index primer, 34*μ*L H_2_O and 10*μ*L of the ExoI-treated cDNA. Samples were cycled using the following program: 1. 95^◦^C 3min 2. 98^◦^C 20sec 3. 67^◦^C 15sec 4. 72^◦^C 15sec. 5. 72^◦^C 1min. Steps 2-4 were repeated for the number of cycles determined by qPCR to prevent overamplification. Libraries were cleaned-up using SPRI beads and eluted in 22*μ*L H_2_O. A second index was added by combining 50*μ*L 2X KAPA HiFi HotStart Ready mix, 2*μ*L 10*μ*M forward index primer, 2*μ*L 10*μ*M reverse index primer, 20*μ*L PCR product (from previous step), 24*μ*L H_2_O. Samples were cycled using the following program: 1. 95^◦^C 3min 2. 98^◦^C 20sec 3. 67^◦^C 15sec 4. 72^◦^C 15sec repeat 2-4 X 8 5. 72^◦^C 1min. Libraries were cleaned-up using SPRI beads, eluted in 22*μ*L H_2_O, and subsequently pooled and size-selected using PAGE. Library size distribution and quantity was assessed using a tapestation instrument (Agilent). Sequencing was performed on a NovaSeqX (Illumina) at the UCSF Center for Advanced Technologies.

### 4.5. Computational analysis of eclair CLIP libraries

Raw paired-end FASTQ files were first converted to single-read ecCLIP input files while preserving molecular barcode information. For each R1 FASTQ file, the matching R2 file was used to append a unique molecular identifier (UMI). Datasets were then analyzed using the CLIP Tool Kit (CTK) based scripts. Unless otherwise noted, reads were trimmed with cutadapt (v2.3) by removing the first 3 nt (-u 3), applying a quality cutoff of 15, retaining reads of at least 20 nt, and trimming a poly(A) adapter sequence (AAAAAAAAAAAAAAAAAA). Trimmed reads were aligned to the human hg38 genome with bwa (v0.7.17) using -n 0.06 and -q 20. Alignments were sorted and indexed with samtools (v1.7), and PCR duplicates were collapsed with umi_tools (v1.1.6) dedup using the UMI encoded in the read name.

Deduplicated alignments were converted to CTK (v1.1.2) tag BED files using parseAlignment.pl with minimum mapping quality 1 and minimum read length 18. Mutation files were generated during parsing, and mutations corresponding to unique tags were retained with selectRow.pl. Biological replicates were then concatenated into pooled tag BED files. CTK tag2profile.pl was used to generate strand-specific bedGraph profiles, and CTK tag2peak.pl was used for peak calling with strand specificity, valley seeking, p = 0.05, valley depth 0.9, hg38 annotation, and multiple-test correction. Both full peak regions, peak boundaries, and half peak-height regions were exported.

Mutations in unique tags were split into substitutions, strand-aware C-to-T conversions, deletions, and insertions. Deletion-based crosslinking-induced mutation sites (CIMS) were called from pooled unique tags with CTK CIMS.pl using -n 10, permutation mode, and cached deletion backgrounds. Sites with FDR ≤ 0.1 were retained for the main RBP analyses, and ±10 nt windows around CIMS sites were generated for sequence and motif analyses.

Background-filtered peak and CIMS sets were generated by removing strand-matched intervals overlapping the tag-only negative-control BED set using intersectBed -s -v (bedtools v2.26.0). These filtered BED files were the primary inputs for the atlas-level summaries.

### 4.6. RBP atlas summaries

Filtered CIMS and peak BED files were collected across RBPs. Genomic-region annotations were assigned with strand-specific intersectBed against hg38 3^′^ UTR, 5^′^ UTR, exon, and intron BED files. Categories were assigned hierarchically: 3^′^ UTR first, then 5^′^ UTR after excluding 3^′^ UTR intervals, then coding/exonic intervals after excluding UTRs, then introns after excluding UTRs and exons. Remaining intervals were classified as intergenic.

For atlas-level overlap analysis, filtered intervals were concatenated, sorted, and merged strand-specifically with mergeBed -s. Merged intervals were named by genomic coordinates and the set of contributing RBPs. Per-RBP binary/count matrices were generated by intersecting each RBP’s filtered BED file with the merged interval set using intersectBed -wo -s, followed by counting overlaps per merged interval. A separate tag-count matrix was produced by intersecting each pooled unique-tag BED file with the merged interval set. Heatmaps and clustered correlation plots were generated with pandas, seaborn, matplotlib, and scipy.

Biological replicate concordance was estimated by converting filtered BED intervals to SAF annotation files and counting deduplicated BAM reads over those intervals with featureCounts (v1.6.2) -F SAF -R BAM -s 1 -T 20. For each RBP, replicate count columns were read into pandas and Pearson correlation coefficients were calculated with scipy.

### 4.7. Sequencing Read Processing

Paired-end sequencing reads were processed separately for each RBP, sample type, and replicate. Read pairs were first merged with PEAR using multithreading. PEAR intermediate FASTQ outputs were compressed after merging. The terminal 8 nt UMI was extracted from the 3^′^ end of each assembled read using umi_tools extract –3prime with the pattern NNNNNNNN. The extracted UMI was appended to the read name and the UMI bases were removed from the read sequence.

Library linkers were removed from UMI-trimmed assembled reads using cutadapt with a linked-adapter model matching the expected 5^′^ and 3^′^ library linkers. Reads were retained only if the expected linker structure was detected. After linker removal, inserts were filtered to retain only exact designed insert lengths: 50 nt for SNP constructs and 75 nt for structure-variant constructs.

### 4.8. Reference Construction and Alignment

Per-RBP reference FASTA files were generated from the construct metadata. Metadata sequences were provided as RNA sequences and were converted to DNA by replacing U with T. SNP references included both reference and alternate 50 nt alleles. Structure references included reference controls, structure-preserving variants, structure-disrupting variants, and negative-control variants. FASTA headers were sanitized to Bowtie-compatible construct identifiers, with a separate metadata table preserving the mapping to the original construct IDs and annotations.

Filtered inserts were aligned to the corresponding per-RBP reference using Bowtie2 in end-to-end mode. Alignments were restricted to exact insert matches by using –score-min L,0,0; in end-to-end alignment this requires a Bowtie2 score of 0. Reads were aligned with –norc, –very-sensitive, -L 18, -k 2, and –no-unal. Only reads with a unique best exact alignment were counted. PCR duplicates were collapsed by counting unique UMIs per construct per sample.

### 4.9. Count Normalization

For each RBP, a construct-by-sample count matrix was generated from unique UMI counts across IP and input samples for both biological replicates. SNP and structure constructs were normalized together because they were present in the same library and sequencing samples.

Counts were normalized with a conservative DESeq2-style median-of-ratios procedure. Size factors were estimated from construct count ratios and then rescaled so that the minimum size factor was 1.0. This ensures that normalization only scales counts downward and does not inflate counts or statistical power. Normalized counts were floored to integers before logistic-regression analyses.

### 4.10. SNP Effect Analysis

For each SNP, reference and alternate allele constructs were compared using a binomial logistic-regression model adapted from the Ribolog framework. For each replicate and allele, IP counts were modeled as successes and input counts as failures. The model included allele and replicate terms:

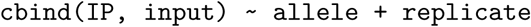

The reference allele was used as the treatment reference level. The allele coefficient therefore estimates the log-odds-ratio of IP/input for the alternate allele relative to the reference allele. This coefficient was converted to log_2_ units:

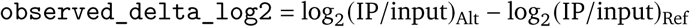

SNP model p-values were obtained from the Wald test for the alternate-allele coefficient. Multiple-testing correction was performed using Benjamini-Hochberg FDR both globally across all tested SNPs and separately within each RBP. Positive predicted values indicate higher predicted RBP binding for the alternate allele relative to the reference allele, while negative values indicate lower predicted binding for the alternate allele.

### 4.11. Structure-Variant Analysis

Structure-variant constructs were grouped by reference peak. Each construct was assigned to one of four analysis groups: Ref (reference), Structure-preserv. (structure preserving mutations), Structure-disrupt. (structure disrupting mutations), Neg. set (negative set; multiple mutations to disrupt both sequence and structure). For each construct, log_2_ enrichment was calculated from normalized IP and input counts using a small pseudocount. Replicates were combined at the construct level for plotting, so each construct contributed a single plotted value. For pairwise group comparisons, binomial logistic regression modeled normalized IP counts as successes and normalized input counts as failures, with construct-group and replicate terms. The construct-group coefficient tests a difference in IP/input enrichment between the two groups after accounting for replicate effects. We required log_2_(IP/input) for Ref to be >0.25, so that we are confident the sequence is being bound by the cognate RBP within the context of the reporter.

Predicted structural and predicted not-structural peaks were compared using effect-size summaries, one-sided Mann-Whitney tests, Welch t-tests, and Cliff’s delta. A categorical expected-pattern analysis was also performed, where a peak was counted as matching the expected structure-dependent pattern when it was reference-enriched, Structure-preserv. was closer to Ref than Structure-disrupt. was, and Structure-preserv. exceeded Structure-disrupt.. Enrichment of this pattern among predicted structural peaks was evaluated with a one-sided Fisher exact test. For the primary paired rescue analysis, we compared absolute distances from reference enrichment for structure-preserving and structure-disrupting constructs within each reference-enriched site using a one-sided paired Wilcoxon test, separately for the two predicted structural classes (**Fig. 4c**).

### 4.12. Pythia model

#### 4.12.1 Overview

Pythia is a dual-arm neural network architecture that jointly processes RNA sequence and potential base-pairing information to predict either RBP binding (binary classification) or RNA structural properties (four BEACON tasks: contact map prediction, distance map prediction, secondary structure prediction, and structural score imputation). All models share a common structural inductive bias: a frozen fixed dilated convolutional layer that explicitly encodes candidate Watson–Crick and wobble base pairs across a range of stem distances and bulge configurations.

#### 4.12.2 Input representation

RNA sequences are encoded as one-hot tensors of shape (B, 4, L), where B is batch size, L is sequence length, and the four channels correspond to A, C, G, and U (T is treated as U). All sequences are zero-padded or truncated to the task-specific maximum length prior to batching.

#### 4.12.3 Fixed dilated convolutional layer

The core structural feature extractor is a single frozen 1-D convolutional layer with pre-designed kernels that score potential base pairs at multiple distances and bulge configurations. For each dilation radius d ∈ [d_ start, d_end] and each Watson–Crick or wobble pair type (A-U, U-A, C-G, G-C, G-U, U-G), the layer places one kernel that activates when a complementary nucleotide pair is present at positions separated by d. To accommodate imperfect stems, each dilation radius is further expanded into bulge variants: for bulge size 1, offsets of {−1, 0, +1} are applied to each position in the pair; for bulge size 2, additional offsets of ±2 are included. Kernel weights are fixed and not updated during training. The total number of output channels depends on the dilation range and bulge size; for the default configuration (d_start=5, d_end=24, bulge_ size=2) this yields 600 channels.

#### 4.12.4 Dual-arm feature extraction

All Pythia models process each input sequence through two parallel convolutional arms:

##### Structure arm (arm1)

receives the output of the fixed dilated convolutional layer and applies three successive 1-D convolutional blocks (kernel sizes 16, 10, 5) with BatchNorm and ReLU activations between each layer. For the RBP model, an adaptive max-pool to 60 positions is inserted before arm1 to produce a fixed-length representation from the variable-length FD output; for BEACON tasks, the FD output is symmetrically padded and center-cropped to preserve sequence length L.

##### Sequence arm (arm2)

receives the raw one-hot input and applies three successive 1-D convolutional blocks (kernel sizes 12, 7, 5) with BatchNorm and ReLU activations.

After both arms, the feature maps are concatenated along the channel dimension. A BatchNorm1d transition layer is applied to the concatenated representation before downstream processing in all BEACON models (the RBP model omits this).

#### 4.12.5 Pythia RBP model

##### Task

Binary classification of ecCLIP peak sequences (Bound) versus dinucleotide-shuffled control sequences (Unbound). The negative class is a computational control, not an experimentally verified set of unbound transcripts.

##### Architecture

Input sequences are 256 nt. The structure arm (arm1) processes fixed dilated convolutional features (d_start=5, d_end=24, bulge_size=2) after adaptive max-pooling to 60 positions (BatchNorm + ReLU), then applies three Conv1d layers with widths (64, 64, 64), kernel sizes (16, 10, 5), and max-pool stride 2 after each layer. The sequence arm (arm2) applies three Conv1d layers directly to one-hot input with widths (128, 64, 64), kernel sizes (12, 7, 5), and max-pool stride 2 after each. Both arms use dropout (p=0.25). The outputs of both arms are concatenated and flattened, then passed through three fully connected layers: Dense(64) → ReLU → Dense(32) → ReLU → Dense(2). Weights are initialized with Kaiming normal (Conv1d), Xavier uniform (Linear), and constant 0.01 bias (Linear).

##### Training

Models are trained with Adam (lr=1×10^−3^, weight_decay=1×10^−4^), batch size 64, for up to 50 epochs with early stopping (patience=5). A linear learning-rate warmup over 2 epochs is followed by cosine decay. Gradient norms are clipped to 1.0. Training uses 16-bit mixed precision. The loss is CrossEntropyLoss with class weights [0.2, 1.0] (Unbound, Bound), giving the positive (Bound) class 5× more influence to compensate for the ∼10:1 negative-to-positive class imbalance within the training datasets. For each of the 67 RBPs, a separate model is trained. Negative (Unbound) examples are dinucleotide-shuffled sequences derived from bound peaks.

### 4.13. Pythia BEACON models

All four BEACON task models share the same dual-arm backbone (FixedDilatedConv d_start=5, d_end=24 unless noted; bulge_size=2; dropout=0.15) and diverge at the fusion and prediction head. For the three 2-D tasks (CMP, DMP, SSP), per-position 1-D features from both arms are lifted into pairwise (L × L) representations via outer product, outer sum, and absolute outer difference operations, then projected into a shared 2-D representation and refined by a 2-D ResNet. All pairwise 2-D outputs are symmetrized. For the 1-D task (SSI), per-position features are refined by a stack of 1-D residual blocks and decoded by a scalar regression head.

### Learned distance encoding

For all four tasks, a learned positional encoding of pair distance |*i*− *j*| is added to the pairwise feature map. This is parameterized as a lookup table over binned distances (dist_enc_dim=64 for all models), allowing the model to learn distance-dependent priors for structural contact formation.

#### 4.13.1 Contact map prediction (PythiaCMP)

##### Task

Binary prediction of long-range RNA base contacts (|*i* − *j*| ≥ 23).

##### Architecture

Arm1 widths: (256, 192, 128); Arm2 widths: (256, 128, 128). Total 1-D feature dimension at arm outputs: 128 + 128 = 256. Pairwise feature channels: outer_prod + outer_sum + outer_diff (3 × 256) + dist_enc (64) = 832. A 1×1 Conv2d (BatchNorm + ReLU) projects pairwise features to hidden_dim=256.

The 2-D ResNet contains 7 residual blocks (BatchNorm + ReLU) with a multi-scale dilation pattern [1, 2, 4, 8, 1, 2, 4] applied in sequence. The output head applies Conv2d(256→128, kernel 3) + BatchNorm + ReLU + Dropout + Conv2d(128→1, kernel 1), followed by symmetrization. Sequences are padded to max_len=1024.

##### Loss and metric

BCEWithLogitsLoss. Evaluation uses the BEACON official Top-L precision: for each RNA of length L, the top-L predicted positions (by sigmoid probability) with score > 0.5 among positions with |*i* − *j*| ≥ 23 are taken, and precision is micro-averaged across the test set.

##### Training

Adam (lr=1×10^−4^, weight_decay=0.01), batch size 1, gradient accumulation over 8 steps (effective batch size 8), up to 50 epochs, early stopping patience=5, linear warmup for 1 epoch.

#### 4.13.2 Distance map prediction (PythiaDMP)

##### Task

Regression of pairwise C3^′^–C3^′^ distances (normalized to [0,1]).

##### Architecture

Arm1 widths: (192, 128, 128); Arm2 widths: (256, 192, 128). Total 1-D feature dimension: 128 + 128 = 256. Pairwise feature channels: outer_prod + outer_sum + outer_diff + distance-weighted outer_diff + inverse-distance-weighted outer_sum (5 × 256) + raw sequence distance (1) + multi-scale exponential decay features at scales 10, 50, 100 nt (3) + dist_enc (64) = 1348. A 1×1 Conv2d (BatchNorm + ReLU) projects to hidden_dim=228. The 2-D ResNet contains 7 blocks with dilation pattern [1, 2, 4, 8, 1, 2, 4]. The output head applies Conv2d(228→114) + BatchNorm + ReLU + Dropout + Conv2d(114→1), followed by sigmoid (to enforce [0,1] output) and symmetrization. Sequences are padded to max_len=1024.

##### Loss and metric

Huber loss (*δ*=0.1). Evaluation uses global Pearson R^2^ computed over all non-padded pairwise positions across the test set (micro-averaged).

##### Training

Adam (lr=5×10^−5^, weight_decay=0.01), batch size 1, gradient accumulation over 8 steps (effective batch size 8), up to 50 epochs, early stopping patience=5, linear warmup for 5 epochs.

#### 4.13.3 Secondary structure prediction (PythiaSSP)

##### Task

Binary prediction of base-paired positions (full L × L contact matrix, including short-range pairs).

##### Architecture

Arm1 widths: (128, 128, 128); Arm2 widths: (256, 128, 128). Total 1-D feature dimension: 128 + 128 = 256. Pairwise feature channels: outer_prod + outer_sum + outer_diff (3 × 256) + dist_enc (64) = 832. A 1×1 Conv2d (GroupNorm(8) + GELU) projects to hidden_dim=256. The 2-D ResNet contains 8 residual blocks (GroupNorm(8) + GELU) with dilation pattern [1, 2, 4, 1, 2, 4, 1, 1]. The output head applies Conv2d(256→128, kernel 3) + GroupNorm(8) + GELU + Conv2d(128→1, kernel 1), followed by symmetrization. Sequences are padded to max_len=512.

##### Loss and metric

BCEWithLogitsLoss. Evaluation uses global micro-averaged F1 (with precision and recall) at a 0.5 sigmoid threshold across all non-padded positions.

##### Training

Adam (lr=3×10^−4^, no weight decay), batch size 4, gradient accumulation over 2 steps (effective batch size 8), up to 100 epochs, early stopping patience=3, linear warmup for 3 epochs.

#### 4.13.4 Structural score imputation (PythiaSSI)

##### Task

Per-position regression of structural accessibility scores (e.g., SHAPE reactivity or predicted unpairedness), imputing masked positions from context.

##### Architecture

Extended dilation range: d_end=48, giving the fixed dilated conv layer sensitivity to potential base pairs up to 48 nt apart. Arm1 widths: (128, 128, 128); Arm2 widths: (256, 128, 128). Combined 1-D feature dimension after concatenation and transition BatchNorm: 256. A 1×1 Conv1d projects to hidden_ dim=256. Six 1-D residual blocks (BatchNorm + ReLU, dilation=1) refine per-position features. A final Layer-Norm is applied before the regression head: Linear(256→128) + GELU + Dropout(0.15) + Linear(128→1), producing a per-position scalar. Sequences are padded to max_len=440.

##### Loss and metric

MSELoss computed only at masked (imputed) positions. Evaluation uses Pearson R^2^ between predicted and true structural scores at masked positions.

##### Training

Adam with two learning-rate groups — backbone (lr_feat=3×10^−4^) and regression head (lr_ head=1.5×10^−3^) — weight_decay=0.01; batch size 64, gradient accumulation over 2 steps; up to 40 epochs, early stopping patience=10, linear warmup for 5 epochs. Gradient norms are clipped to 1.0.

### 4.14. DeepLIFT analysis of sequence versus structure contributions

To quantify the relative contributions of sequence and structural features to individual RBP binding predictions, we applied DeepLIFT to models trained on each RBP included in the attribution analysis. For each peak sequence, DeepLIFT attribution scores were computed separately by replacing the sequence arm input or the structure arm input with an all-zero (no-nucleotide) baseline, holding the other arm fixed. The resulting attribution scores reflect the contribution of each arm to the binding prediction. For each sequence, the log_2_ ratio of the sequence-arm attribution to the structure-arm attribution was computed as log_2_(DeepLIFT_seq_ randomized / DeepLIFT_str_randomized). Positive values indicate sequence dominance; negative values indicate structural dominance. To focus on the most informative peaks, only the top 10% of peaks ranked by variance across the three scores (structure, sequence, DeepLIFT) were retained per RBP. The resulting distribution of log_2_ ratios per RBP was tested against a median of zero using a one-sample Wilcoxon signed-rank test.

P-values were corrected for multiple testing using the Benjamini–Hochberg method. RBPs with |median log_2_ ratio| > 0.05 and FDR < 10% were classified as sequence-dominant (positive median) or structure-dominant (negative median); the remaining models were labeled mixed. For the reporter validation in **Supplementary Fig. S4b**, this per-site log_2_ score ratio is termed the Structural Reliance Index (SRI) and we defined predicted structure-dependent sites by SRI < −0.1.

### 4.15. Protein-embedding comparison with ecCLIP binding preferences

Canonical sequences for the 67 RBPs were retrieved from UniProt and embedded with ESM2 model (esm2_ t33_650M_UR50D; Lin et al., 2023). Each protein was represented by the layer-30 embedding averaged over its annotated RNA-binding-domain residues (UniProt domain annotations; full-length protein for the four RBPs without an annotated domain) (**Fig. 1c,d**), or over all residues for the whole-protein comparison (**Supp. Fig. S1i,j**). Binding preferences were demonstrated as the log enrichment of each of the 1,024 5-mers in 20-nt windows centered on ecCLIP peak summits relative to flanking background windows, using up to 100,000 peaks per RBP. After standardizing features within each space, concordance was measured with the modified RV coefficient, a Mantel test on pairwise cosine distances (Spearman *ρ*), and distance correlation (reported as R). Significance was assessed against a null from 9,999 permutations of RBP labels. Dendrograms were built with Ward’s method and cut at k = 6, and both trees were colored by the protein-embedding clusters.

### 4.16. Linear probing for RBP prediction

To validate Pythia on RBP-binding prediction, we benchmarked it against large pre-trained RNA foundation models, where we evaluated six RNA language models by linear probing: RNA-FM (Chen et al., 2022), Ri-NALMo (Penić et al., 2025), RNA-MSM (Zhang et al., 2024), RNAErnie (Wang et al., 2024), and the two UTR-LM variants UTR-LM-MRL and UTR-LM-TE_EL (Chu et al., 2024). All models were accessed as pretrained weights through the MultiMolecule framework (Chen, 2026), an open-source library that distributes RNA foundation models via the Hugging Face Hub and were kept frozen throughout. For each of the 67 RBPs, every RNA sequence was tokenized and passed through the frozen model, and a fixed-length representation was obtained by mean-pooling the final hidden-layer token embeddings over all non-special, non-padding positions (“mean-last” pooling). On top of these embeddings, we trained an L2-regularized logistic-regression probe (scikit-learn (Pedregosa et al., 2011); lbfgs solver, class-balanced weighting, up to 1,000-2,000 iterations) to classify bound versus unbound sequences. The inverse regularization strength C was selected per RBP and per model over a grid spanning 10^−3^ to 10^2^, choosing the value that maximized area under the receiveroperating-characteristic curve (auROC) on the held-out tuning set; no foundation-model weights were updated at any stage. Probes were then evaluated on the validation and held-out sets using auROC and area under the precision-recall curve (auPR).

To ensure a controlled comparison, all foundation-model probes and Pythia were trained, tuned, and evaluated on identical per-RBP data splits (training, tuning, validation, and held-out sets) drawn from the eclair-CLIP dataset, with unbound (negative) sequences generated by dinucleotide-preserving shuffling of bound peaks, matching the Pythia RBP training procedure. Using the same splits, we additionally trained five supervised baselines from scratch on each RBP: RBPNet (Her et al., 2026), DeepBind (Alipanahi et al., 2015), DeepSEA (Zhou and Troyanskaya, 2015), PRIESSTESS (Laverty et al., 2022), and GraphProt (Maticzka et al., 2014), evaluated with the same auROC and auPR metrics. For **Fig.** 3**d**,**e**, models were each trained with three random seeds, and per-RBP auROC and auPR were averaged across seeds; linear probes on frozen language-model embeddings are deterministic.

To measure precision in the high recall region, we computed average precision over the recall range 0.9–1 (false-negative rate ≤ 10%) and divided it by the width of this range (0.1), so that a perfect classifier scores 1. For the training-data efficiency analysis (**Supp. Fig. S3g**), we subsampled the training sets of the 12 RBPs in **Fig. 3d** to 10%, 25% and 50%, stratified by label, with three subsamples at each fraction. All six supervised models were retrained on the same subsets, and the tuning and validation sets were left unchanged. DeepSEA and DeepBind kept their original architectures and training procedures. The one exception was DeepSEA, which did not converge with its default settings, so we trained it with a learning rate of 10^−4^ and no weight decay. GraphProt and PRIESSTESS were run with default parameters. All models were scored on the same 10,000 validation sequences.

### 4.17. BEACON Benchmark Suite

To place Pythia’s structure-awareness on a standardized external footing, we also evaluated it on the BEACON RNA-benchmark suite (Ren et al., 2024), focusing on its four structure-oriented tasks: contact map prediction (CMP), distance map prediction (DMP), structural score imputation (SSI), and secondary structure prediction (SSP). For each task we used BEACON’s provided datasets, train/validation/test splits, and evaluation protocol without modification, so that Pythia’s results are directly comparable to the 17 baseline models reported in the benchmark; three conventional architectures (CNN, ResNet, LSTM), twelve pre-trained RNA language models (RNA-FM, RNABERT, RNA-MSM, three SpliceBERT and four 3UTRBERT variants, and two UTR-LM variants), and the benchmark’s own BEACON-B and BEACON-B512 models. The Pythia task-specific architectures and training configurations are described above (Pythia BEACON models); in every case only the task-specific arms and prediction head were trained while the fixed dilated-convolution layer remained frozen. For each task the best-performing epoch on the validation split was selected and evaluated once on the held-out test set. Task-specific metrics followed the BEACON definitions: CMP was scored by Top-L precision over long-range contacts (contacts defined at an 8 Å threshold), DMP by the coefficient of determination (R^2^) over valid pairwise positions, SSI by R^2^ over masked positions, and SSP by F1 over base-paired positions.

### 4.18. Secondary-structure accessibility of the HNRNPC poly-U binding site

HNRNPC recognizes poly-U only when the tract is single-stranded and presented across the RRM *β*-sheet (PDB 2MXY; Cieniková et al., 2014; Cienikova et al., 2015), so we used RNA secondary structures to test whether the structure-preserving and structure-disrupting mutations remain compatible with binding. Minimum-free-energy secondary structures for each 100-nt construct were predicted from sequence with ViennaRNA (RNAfold v2.7.2, default Turner 2004 parameters; Lorenz et al., 2011), and per-nucleotide base-pairing probabilities were derived from the equilibrium partition function. Structures were drawn with VARNA (Darty et al., 2009) and accessibility was quantified as the number of U52-U58 bases left unpaired in the minimum-free-energy structure and their mean probability of being unpaired.

### 4.19. Deep Learning-based joint modeling of RBP binding, mRNA degradation and expression

#### 4.19.1 Model architecture

The Borzuki model follows the same overall architecture as the original Borzoi model (Linder et al., 2025). However, since we trained Borzuki on a considerably smaller set of coverage profiles, we reduced the number of parameters. Specifically, we trained two different model sizes, each with a fixed input window size of 393,216 bp:

40 million parameters:

- Convolution tower: Starts with 320 channels, ends with 768 channels.
- Self-attention: 8 blocks, 64 key size, 2x dense expansion.

5 million parameters:

- Convolution tower: Starts with 192 channels, ends with 384 channels.
- Self-attention: 4 blocks, 32 key size, 1.5x dense expansion.

#### 4.19.2 Training data

We curated RNA-seq degradation time-course experiments from recount3, and paired these data with auxiliary post-transcriptional and epigenomic assays from Borzoi’s original training set. These data include:

RNA-seq degradation time-course experiments:

- 14 experiments in human HEK293T, HeLa, MCF7, HepG2, LCL, Jurkat, K562, and HUES9 cells.
- 11 experiments in mouse 3T3, mESC, cortical neuron, and dendritic cells.

CLIP-seq experiments:

- 31 ENCODE eCLIP experiments in human K562 and HepG2 cells.
- 11 ecCLIP-seq experiments from the Goodarzi Lab in HEK293T cells.

Perturb-seq experiments:

- 10 perturbations of 3^′^-cleavage regulators in HEK293T cells (Kowalski et al., 2024).

Baseline training data from the original Borzoi model:

- FANTOM5 CAGE, ENCODE DNase-seq, and RNA-seq matching any of the above 12 cell types.

See the ‘Data and code availability’ section for access to a detailed list of data identifiers. We only included CLIP-seq targets that overlapped the set of stability-correlated RBPs identified by Agarwal and Kelley (2022).

#### 4.19.3 Marginalized variant effect predictions

To make predictions for genetic variants measured in an MPRA context, we inserted the reference and alternate oligo sequences into multiple native genomic loci and averaged the variant effect score across these contexts. The model makes variant effect predictions for multiple RNA-seq tracks, and these values must be distilled into a scalar variant effect score. We tested two methods for making such marginalized predictions, which are detailed below.

As preliminaries, let ***s***^ref^ and ***s***^alt^ be the reference and alternate oligos. Insert ***s***^ref^ and ***s***^alt^ into *K* distinct genomic sequences (each representing a unique gene’s 3^′^ UTR), producing a set of reference and alternate one-hot-coded sequences 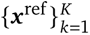 and 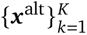, each of shape *L* × 4 (*L* = 393,216 for Borzuki). Obtain coverage predictions ***y***^ref^ , ***y***^alt^ ∈ ℝ*^M^*^×*K*×*T*^ for all *T* RNA-seq tracks (*M* = output length) and compute exon-aggregated log-coverage statistics ***u***^ref^ , ***u***^alt^ ∈ ℝ*^K^*^×*T*^ :

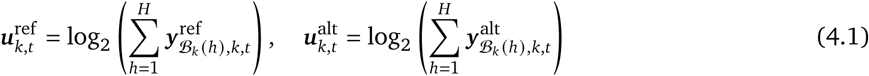

The set B*_k_* = {*b*_1_, . . . , *b_H_* } denotes the indices of exon-overlapping output bins for the target gene context *k*.

#### 4.19.4 Approach 1 -Fit to Saluki PC1

In this method, we fit a linear regression model to predict Saluki’s cell type-agnostic half-life measurements, which were originally derived from a principal component analysis of multiple uniformly processed RNA degradation datasets (Agarwal and Kelley, 2022). The idea is to use this regressor as a heuristic to distill scalar half-life variant effect predictions from Borzuki’s many output tracks. First, we assume access to Borzuki’s predicted exon-aggregated log-coverage statistics for all *N* human genes of the original training dataset ***u*** ∈ ℝ*^N^*^×*T*^. We fit a linear regression model *f* to predict Saluki’s PC1 half-life measurements ***v*** ∈ ℝ*^N^* given ***u*** as input features. Using this model, we can predict a scalar effect *z* ∈ ℝ for a given MPRA variant as:

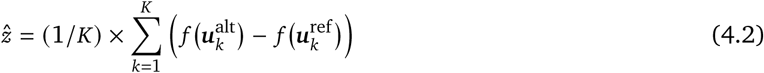

In other words, the variant scores derived from each RNA-seq track *t* are scaled by the regression weight ***w****_t_* corresponding to that track’s importance for predicting the Saluki measurement, averaged across *K* contexts.

#### 4.19.5 Approach 2 -Fit to MPRA reference values

This method is simpler than Approach 1, but requires access to individual MPRA measurements of the reference oligo sequences. To this end, let ***v***^ref^ ∈ ℝ*^N^* be the log-transformed MPRA reference values (e.g. reference oligo log-decay rates), and let ***U***^ref^ ∈ ℝ*^K^*^×*N*×*T*^ be the log-coverage predictions computed for all *N* oligos. For a given genomic context *k*, the oligos are all inserted into the same 3^′^ UTR. As a result, the columns in 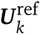 should correlate with ***v***^ref^ , although they will be offset by a constant specific to the particular gene *k* into which we inserted the oligos. Hence, for each context *k*, we fit a linear regression model *f_k_* to predict ***v***^ref^ using 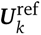 as input features:

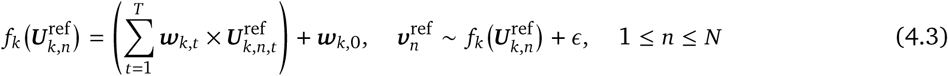

This model is more flexible than Approach 1, as it learns a specific weight ***w****_k_*_,*t*_ for a given context *k* and RNA-seq output track *t*. The model might learn to rely on a particularly salient cell type (e.g. K562) for gene context *k*, while for another context *j* it might rely on different cell types (for example, if context *j* is not expressed in K562 cells).

Finally, to make variant effect prediction *z* ∈ ℝ for a given variant, we calculate the difference in weighted log-coverage, averaged across the *K* linear models:

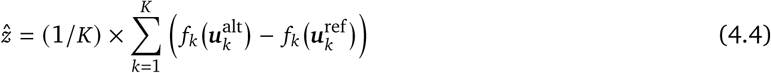

For ensemble predictions, we re-compute and average the predicted effect *z* across multiple Borzuki replicates.

## Supplementary Material

### A. Acknowledgements

We thank A. Pawluk and C. Ricci-Tam for reading earlier versions of this manuscript and helping us with the content. We thank Albertas Navickas for input on eclair CLIP method development.

### B. Funding

L.F. was a Zena Werb Scholar supported by T32CA108462. H.K. was a Zena Werb Scholar supported by T32CA108462. Sequencing was performed at the UCSF Center for Advanced Technology (CAT), supported by UCSF PBBR, RRP IMIA, and NIH 1S10OD028511-01 grants. H.S.N. is supported by the Canadian Institutes of Health Research (CIHR) grant PJT-173317 and resource allocations from Digital Research Alliance of Canada, and holds a CIHR Canada Research Chair. E.M.S. and M.D.L. are supported by Biohub. H.G. is a Core Investigator at Arc Institute, and this research was supported by Arc.

### C. Data, code, and materials availability

All data and code needed to evaluate and reproduce the results in the paper are present in the paper and/or the Supplementary Materials.

- Pythia codebase is available at https://github.com/goodarzilab/pythia.
- Pre-trained Borzuki model weights and code are available at https://github.com/calico/ borzoi-paper/tree/main/extensions/borzuki.
- All raw and processed sequencing data are available from the Gene Expression Omnibus (GEO) under accession number GSE349934.

### D. Supplementary figures

**Figure S1.**
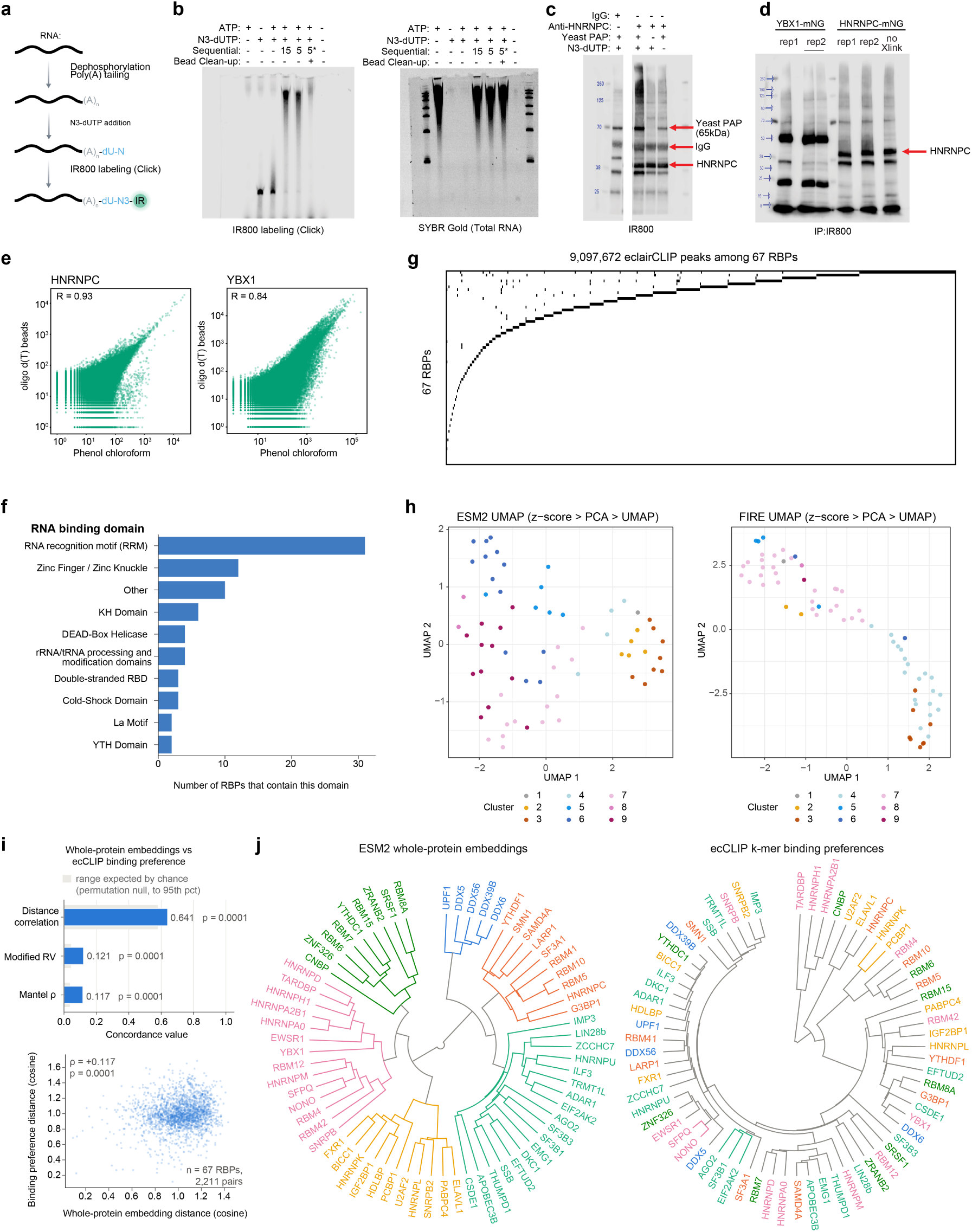
Optimization and validation of the eclair CLIP workflow. (a) Schematic of eclair CLIP RNA labeling strategy. **(b)** An RNA oligo was subjected to poly(A) tailing with Yeast PAP, N_3_-dUTP azide incorporation, and IR800 click labeling. Denaturing PAGE comparing labeling efficiency across conditions varying ATP, N_3_-dUTP concentration, sequence and timing of steps, and bead cleanup, visualized by IR800 (left) and SYBR Gold stain (right). Sequential (-) indicates co-addition of ATP and N_3_-dUTP. Sequential (+) indicates ATP addition followed by N_3_-dUTP addition with the number indicating time in minutes of N_3_-dUTP incubation. Sequential* indicates ATP addition followed by SPRI clean-up followed by N_3_-dUTP incubation for 5 minutes. **(c)** Blot showing immunoprecipitation of HNRNPC from UV-crosslinked HEK293T cells using an anti-HNRNPC antibody or anti-mouse IgG. Shown are IgG control compared to anti-HNRNPC pulldown with and without yeast PAP-mediated poly(A) tailing and N_3_-dUTP labeling. All labeled with DBCO-IR800. Bands corresponding to yeast PAP (65 kDa), IgG, and HNRNPC are indicated. **(d)** Blot showing immunoprecipitation of split mNeonGreen-tagged RBPs (YBX1 and HNRNPC) from HEK293T cells. A neon green nanobody was used to immunoprecipitate and the ecCLIP workflow used to label the complexes. A no-crosslink control is included. **(e)** Comparison of RNA isolation methods for eclair CLIP. Scatter plots show per-gene read counts for RNA isolated by oligo d(T) magnetic beads versus phenol chloroform extraction for HNRNPC (Pearson R = 0.93) and YBX1 (Pearson R = 0.84), demonstrating high concordance between methods. **(f)** Distribution of RNA binding domains among proteins in the ecCLIP dataset. Some proteins contain multiple RNA binding domain types. **(g)** Distribution of ecCLIP peaks across 67 RBPs. Dot plot shows the number of peaks called per RBP, ranked by peak count. A total of 9,097,672 peaks were identified across all 67 RBPs. **(h)** UMAP projections of ESM2 protein embeddings (left) and ecCLIP-derived sequence preferences labeled FIRE (right), following z-score normalization and principal component analysis. Points represent RBPs and are colored by clusters within each representation. **(i)** Whole-protein version of the analysis in Fig. 1c: concordance statistics between ESM2 whole-protein embeddings and ecCLIP-derived binding preferences, shown against the permutation-null range (grey bands), with the corresponding pairwise cosine-distance scatter (2,211 RBP pairs). **( j)** Circular dendrograms as in Fig. 1d, built from whole-protein embeddings (left) and ecCLIP-derived binding preferences (right), both colored by the protein-side grouping.

**Figure S2.**
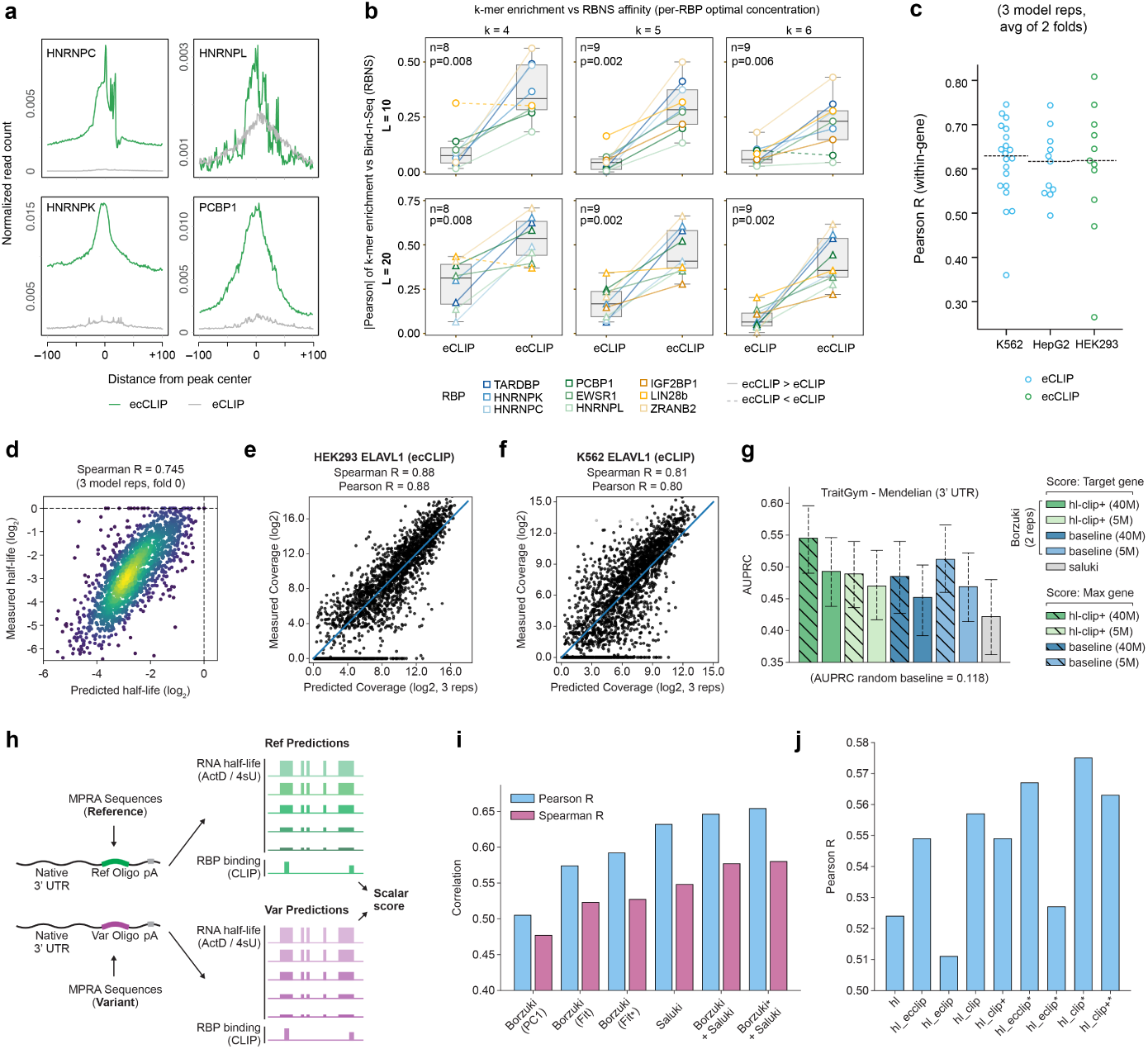
Extended ecCLIP quality metrics and Borzuki validation. **(a)** Additional aggregate profiles comparing ecCLIP and eCLIP read signal centered on peak summits for matched RBPs. **(b)** Robustness of k-mer concordance to RBNS concentration choice. The absolute Pearson correlation between CLIP-derived k-mer enrichment and RBNS affinity is shown using the optimal RBNS protein concentration for each RBP, rather than the fixed 1300 nM concentration used in Fig. 2c. Results are shown for k = 4, 5, and 6 and peak-centered windows of 10 or 20 nucleotides. Lines connect matched RBPs between eCLIP and ecCLIP; solid lines indicate stronger concordance for ecCLIP and dashed lines indicate stronger concordance for eCLIP. P-values correspond to paired comparisons between eCLIP and ecCLIP. **(c)** Test set Pearson R when predicting CLIP-seq profiles at bin-level resolution within genes for held-out test data, shown for both ecCLIP and ENCODE eCLIP tracks. **(d)** Predicted versus measured mRNA degradation rates derived from a K562 actinomycin D transcriptional shutoff experiment. Predicted half-lives were derived by fitting exponential decay curves to Borzuki-predicted exon-aggregated coverage across time points. Spearman R = 0.745 with experimental measurements; Spearman R = 0.728 with Saluki’s cell type-agnostic PC1 half-life summary. **(e)** Predicted versus measured gene-level exon-aggregated log-coverage for the ELAVL1 CLIP-seq experiment in the ecCLIP dataset. **(f)** Predicted versus measured gene-level exon-aggregated log-coverage for an ENCODE eCLIP ELAVL1 experiment. **(g)** Comparison between Saluki and Borzuki when classifying putative causal 3^′^ UTR variants associated with Mendelian traits from TraitGym (Benegas et al., 2025). Variant scoring is compared using the annotated target gene or the gene context with the largest predicted effect. **(h)** Illustration of variant scoring procedure when predicting mRNA half-life effects measured in an MPRA experiment in Beas2B cells (Siegel et al., 2022). Reference and variant oligos are inserted in endogenous 3^′^ UTRs and their predicted effect on RNA half-life and CLIP binding is calculated using Borzuki. For each RNA-seq or CLIP-seq track predicted by the model, the total log fold-change in coverage across the gene’s exons due to the mutation is calculated. These functional multi-track scores are combined into a scalar variant impact using a secondary regressor model, which is fitted either on reference measurements from the MPRA assay or PCA values from the Saluki model’s training data (Methods). **(i)** Comparison between Saluki and Borzuki at the task of predicting half-life MPRA variant effects (’PC1’ / ’Fit’ = Borzuki’s multi-track scores are either fit to Saluki’s PC1 measurements or to the MPRA’s reference values, ‘*’ = variant scores are derived from CLIP-seq tracks in addition to half-life RNA-seq tracks). **( j)** Correlation metrics when predicting MPRA-derived half-life variant effects using versions of Borzuki models trained on subsets of training data (‘hl’ = trained on base ENCODE/FANTOM5 DNase/RNA-seq/CAGE tracks and RNA half-life time-course tracks, ‘hl_ecclip’ = additionally trained on ecCLIP experiments, ‘hl_eclip’ = additionally trained on ENCODE eCLIP, ‘hl_clip’ = trained on ecCLIP and eCLIP tracks, ‘hl_clip+’ = trained on all CLIP-seq experiments and polyadenylation Perturb-seq. ‘*’ = variant scores are derived from CLIP-seq tracks in addition to half-life RNA-seq tracks).

**Figure S3.**
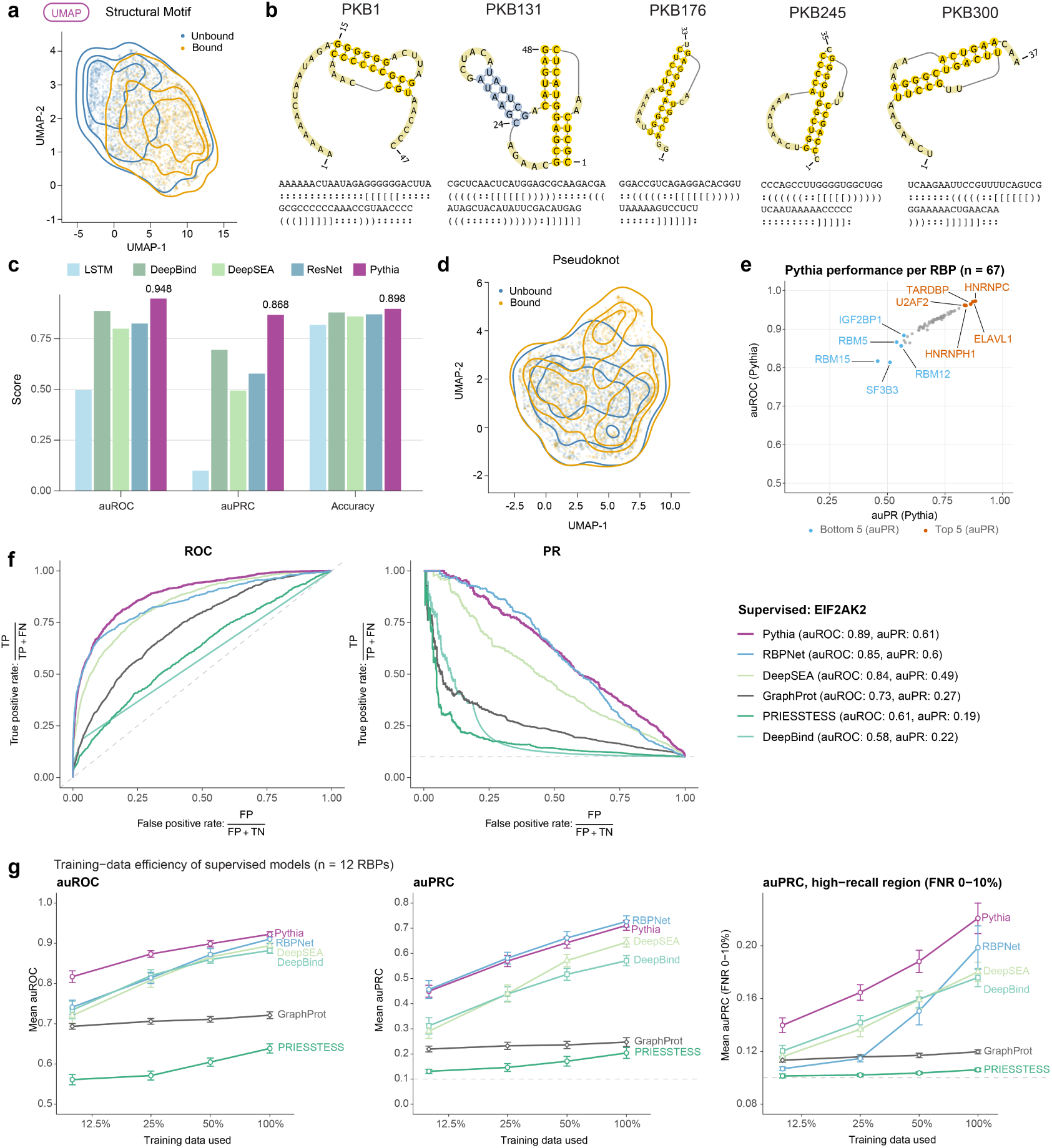
Performance of AI models in predicting RBP binding. **(a)** Generalization to sequence-independent motifs. UMAP projection of Pythia embeddings for held-out synthetic sequences with or without the CFG-defined structural motif. The two groups overlap; this visualization alone does not establish motif classification accuracy. **(b)** Pseudoknot structure prediction. We performed in silico mutations on five distinct pseudoknot structures, generating positive examples that preserved base-pairings and negative examples that disrupted them. **(c)** Pythia distinguished preserved from disrupted pseudoknot structures with an AUROC = 0.95 and AUPRC = 0.87, outperforming other models (LSTM, DeepBind, DeepSEA, ResNet) and demonstrating structural sensitivity. **(d)** Latent space organization. UMAP projection of Pythia embeddings for preserved and disrupted pseudoknot examples. The overlapping distributions are shown descriptively; classification performance is quantified in (c). **(e)** Scatterplot of Pythia auPR and auROC performance per RBP. **(f)** ROC (left) and precision-recall (PR) curves for supervised models predicting genome-wide binding of EIF2AK2. The y-axis of the ROC plot shows true positive rate [TP/(TP+FN)] and the x-axis shows false positive rate [FP/(FP+TN)]. The y-axis of the PR plot shows precision [TP/(TP+FP)] and the x-axis shows recall [TP/(TP+FN)]. Pythia (purple; auROC = 0.89, auPR = 0.61) outperforms RBPNet (auROC = 0.85, auPR = 0.60), DeepSEA (auROC = 0.84, auPR = 0.49), GraphProt (auROC = 0.73, auPR = 0.27), PRIESSTESS (auROC = 0.61, auPR = 0.19), and DeepBind (auROC = 0.58, auPR = 0.22). **(g)** Training-data efficiency of supervised models. Models were trained on 10, 25, 50 or 100% of each RBP’s training set (stratified subsamples, identical across models; three seeds) for the 12 RBPs in Fig. 3d and evaluated on the same 10,000 validation sequences. Left, mean auROC; middle, mean auPRC; right, mean auPRC in the high-recall region (recall 0.9–1). Error bars, s.e.m. across RBPs.

**Figure S4.**
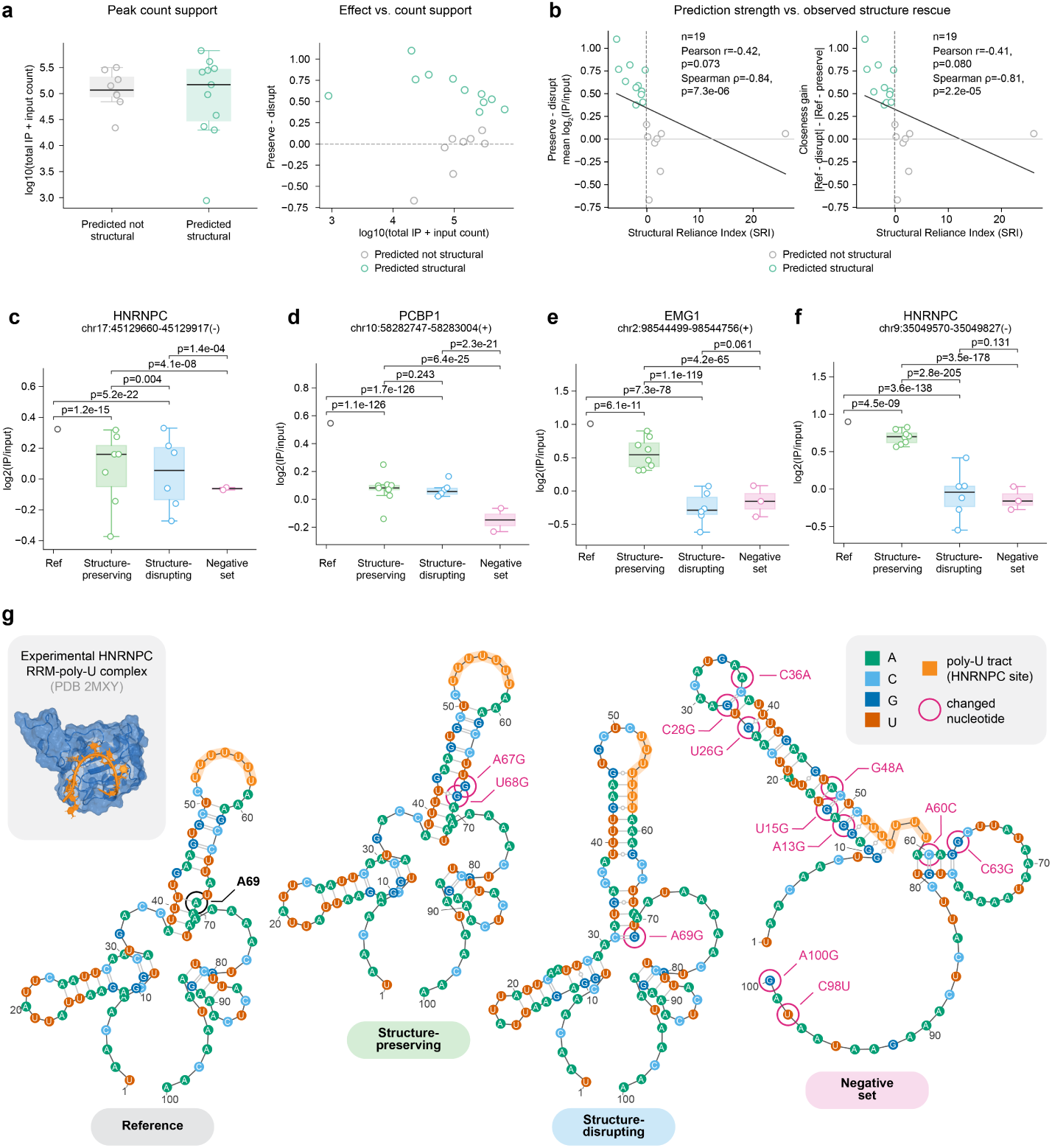
Extended MPRC support for Pythia-nominated structure-dependent RBP binding. **(a)** Count support for structure-rescue summaries. Each point is one reference-enriched peak with Ref log_2_(IP/input) > 0.25. The left panel compares total peak counts, computed as summed IP plus input counts across constructs, between predicted not-structure-dependent and predicted structure-dependent groups on a log_10_ scale. The right panel plots the preserve-minus-disrupt rescue metric against total peak count to test whether observed rescue effects are concentrated in low-or high-count peaks. Points are colored by predicted structural class. **(b)** Pythia prediction strength versus observed structure-rescue metrics. Each point is one reference-enriched peak with Ref log_2_(IP/input) > 0.25. The x-axis shows a Structural Reliance Index (SRI), defined per peak as log ratio of sequence-arm to structure-arm DeepLIFT attribution under a zero-baseline reference, and the dashed vertical line marks the prediction threshold of −0.1 used to define predicted structure-dependent peaks. The left panel shows preserve-minus-disrupt, defined as mean structure-preserving enrichment minus mean structure-disrupting enrichment. The right panel shows closeness gain, defined as abs(Ref -Structure-disrupting) -abs(Ref -Structure-preserving). Positive y-values indicate a stronger structure-preserving rescue pattern. Points are colored by threshold-defined prediction group; fitted lines and Pearson/Spearman statistics summarize continuous-score associations. **(c–f)** Additional representative structure-variant validation examples for HNRNPC at chr17:45,129,660-45,129,917(-), PCBP1 at chr10:58,282,747-58,283,004(+), EMG1 at chr2:98,544,499-98,544,756(+), and HNRNPC at chr9:35,049,570-35,049,827(-). Each point is one construct summarized across replicates as log_2_(IP/input). Constructs are grouped as reference, structure-preserving, structure-disrupting, or negative-control reporters. Boxes are shown for groups with multiple constructs, overlaid open points show individual construct values, and brackets report pairwise binomial logistic-regression P-values where valid. **(g)** Structural context for the HNRNPC validation example. The experimentally determined HNRNPC RRM-poly-U complex (PDB 2MXY; Cieniková et al., 2014; Cienikova et al., 2015) is shown alongside sequence-based secondary-structure predictions for reference, structure-preserving, structure-disrupting, and negative-control constructs. The poly-U tract is highlighted in orange and altered nucleotides are circled. RNA structures were predicted with ViennaRNA and displayed with VARNA (Methods; Lorenz et al., 2011; Darty et al., 2009). These predictions provide structural context and are not direct measurements of reporter folding in cells.

**Figure S5.**
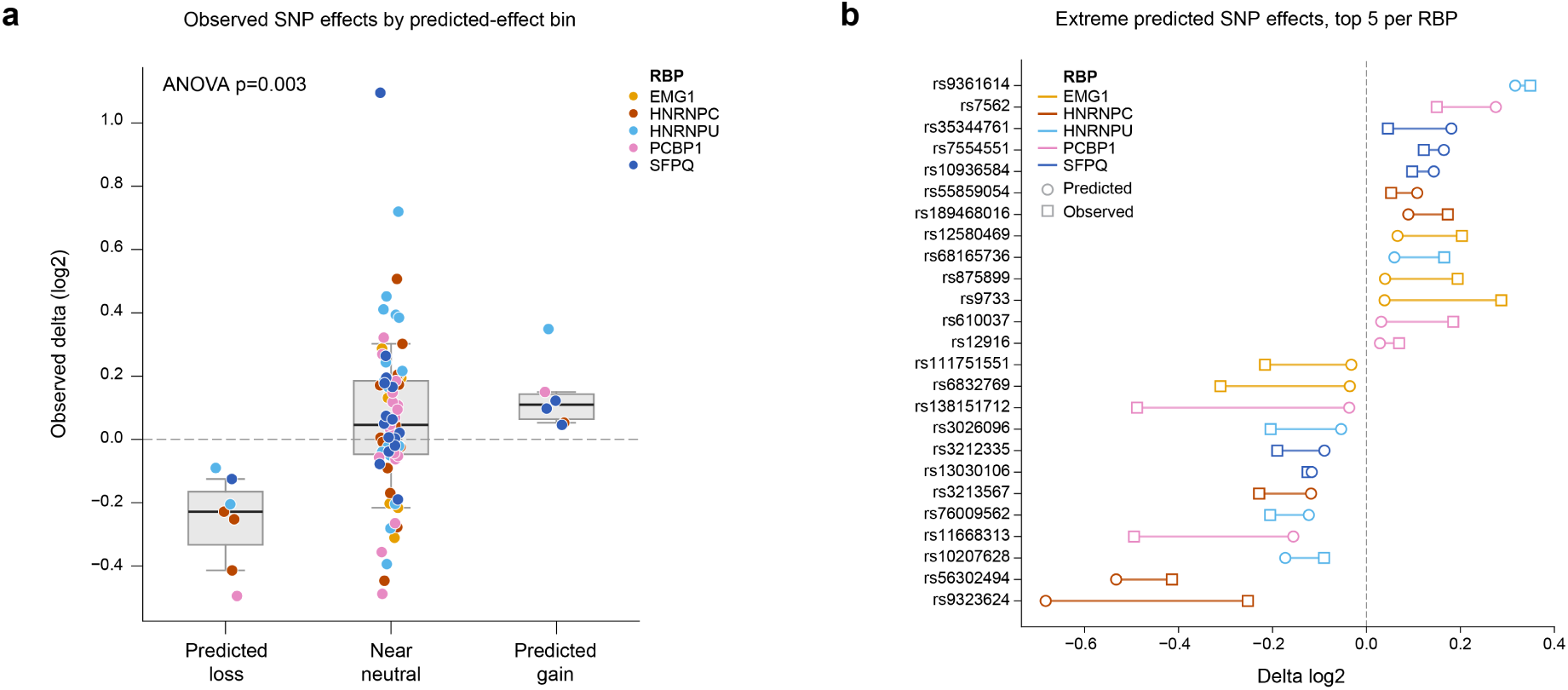
Predicted-effect stratification and high-leverage SNPs for RBP-specific Pythia variant-effect models. **(a)** Observed SNP effects grouped by predicted-effect bin. SNPs with valid, converged model fits were binned by Pythia predicted change into predicted loss (delta < −0.1), near neutral (−0.1 ≤ delta ≤ 0.1), and predicted gain (delta > 0.1). The y-axis shows the observed logistic-regression SNP effect in log_2_ units. Boxes summarize each bin, with the box spanning the 25th to 75th percentiles, the median line marking the 50th percentile, and whiskers set to the 10th and 90th percentiles. Overlaid points show individual SNPs colored by RBP. The dashed horizontal line marks zero observed effect. The text box reports the one-way ANOVA P-value across the three observed-delta distributions. **(b)** Extreme predicted SNP effects. For each RBP, the top five SNPs by absolute Pythia v3 predicted effect are shown. Each horizontal segment connects the predicted SNP effect (circle) to the observed logistic-regression effect (square), both in log_2_ units. The y-axis shows SNP names, and segments and markers are colored by RBP. The vertical dashed line marks zero effect. This plot highlights high-leverage predicted SNPs and shows whether the observed direction and magnitude agree with the prediction.

